# Prominent astrocytic alpha-synuclein pathology with unique post-translational modification signatures unveiled across Lewy body disorders

**DOI:** 10.1101/2022.05.29.493876

**Authors:** Melek Firat Altay, Alan King Lun Liu, Janice L. Holton, Laura Parkkinen, Hilal A. Lashuel

## Abstract

Alpha-synuclein (aSyn) is a pre-synaptic monomeric protein that can form aggregates in neurons in Parkinson’s disease (PD), Parkinson’s disease with dementia (PDD) and dementia with Lewy bodies (DLB), and in oligodendrocytes in multiple system atrophy (MSA). Although the accumulation of aSyn in astrocytes has previously been described in PD, PDD and DLB, the biochemical properties of aSyn in this type of pathology and its topographical distribution have not been studied in detail. Here, we present a systematic investigation of aSyn astrocytic pathology, using an expanded toolset of antibodies covering the entire sequence and known post-translational modifications (PTMs) of aSyn in Lewy body (LB) disorders, including sporadic PD, PDD, DLB, familial PD with *SNCA* G51D mutation and *SNCA* duplication, and in MSA. Astrocytic aSyn was mainly detected in the limbic cortical regions of LB disorders, but were absent in key pathological regions of MSA. These astrocytic aSyn accumulations were detected only with aSyn antibodies against the mid N-terminal and non-amyloid component (NAC) regions covering aSyn residues 34-99. The astroglial accumulations were negative to canonical aSyn aggregation markers, including p62, ubiquitin and aSyn pS129, but positive for phosphorylated and nitrated forms of aSyn at Tyrosine 39 (Y39), and mostly not resistant to proteinase K. Our findings suggest that astrocytic aSyn accumulations are a major part of aSyn pathology in LB disorders, and possess a distinct sequence and PTM signature that is characterized by both N- and C-terminal truncations and modifications at Y39. To the best of our knowledge, this is the first description of aSyn accumulation made solely from N- and C-terminally cleaved aSyn species and the first report demonstrating that astrocytic aSyn exists as a mixture of Y39 phosphorylated and nitrated species. These observations underscore the critical importance of systematic characterization of aSyn accumulation in different cell types as a necessary step to capturing the diversity of aSyn species and pathology in the brain. Our findings combined with further studies on the role of astrocytic pathology in the progression of LB disorders can pave the way towards identifying novel disease mechanisms and therapeutic targets.

## INTRODUCTION

Parkinson’s disease (PD), Parkinson’s disease with dementia (PDD), dementia with Lewy bodies (DLB), and multiple system atrophy (MSA) are age-related neurodegenerative diseases that are characterized by the abnormal accumulation and aggregation of alpha-synuclein (aSyn). Hence, they are collectively referred to as synucleinopathies. PD, PDD and DLB typically involve the accumulation of aSyn in the neuronal soma and processes as Lewy bodies (LBs) and Lewy neurites (LNs), respectively (Baba et al., 1998; Spillantini et al., 1998). MSA, on the other hand, is characterized by the inclusion formation mainly in the oligodendrocytes called glial cytoplasmic inclusions (GCIs) (Spillantini et al., 1998a).

In the early 2000s, several studies reported the presence of aSyn-positive astrocytes in LB diseases (Hishikawa et al., 2001; Shoji et al., 2000; Takeda et al., 2000; Terada et al., 2003; Wakabayashi et al., 2000) (Table 1). Yet, despite the increasing number of publications aimed at dissecting the molecular and structural features of aSyn pathology, very little is known about the biochemical properties and distribution of aSyn species associated with the astrocytes, how they form, and what role they may have in the pathogenesis of PD and other synucleinopathies. Only a small number of studies (13) have been published, using a limited number of antibodies, on the characterization of aSyn astrocytic pathology and its relationship to aSyn neuronal pathology, disease stage or duration (Braak et al., 2007; Fathy et al., 2019; Hishikawa et al., 2001; Kovacs et al., 2014, 2012; Nakamura et al., 2016; Shoji et al., 2000; Song et al., 2009; Sorrentino et al., 2019; Takeda et al., 2000; Terada et al., 2003, 2000; Wakabayashi et al., 2000).

**Table 1:**
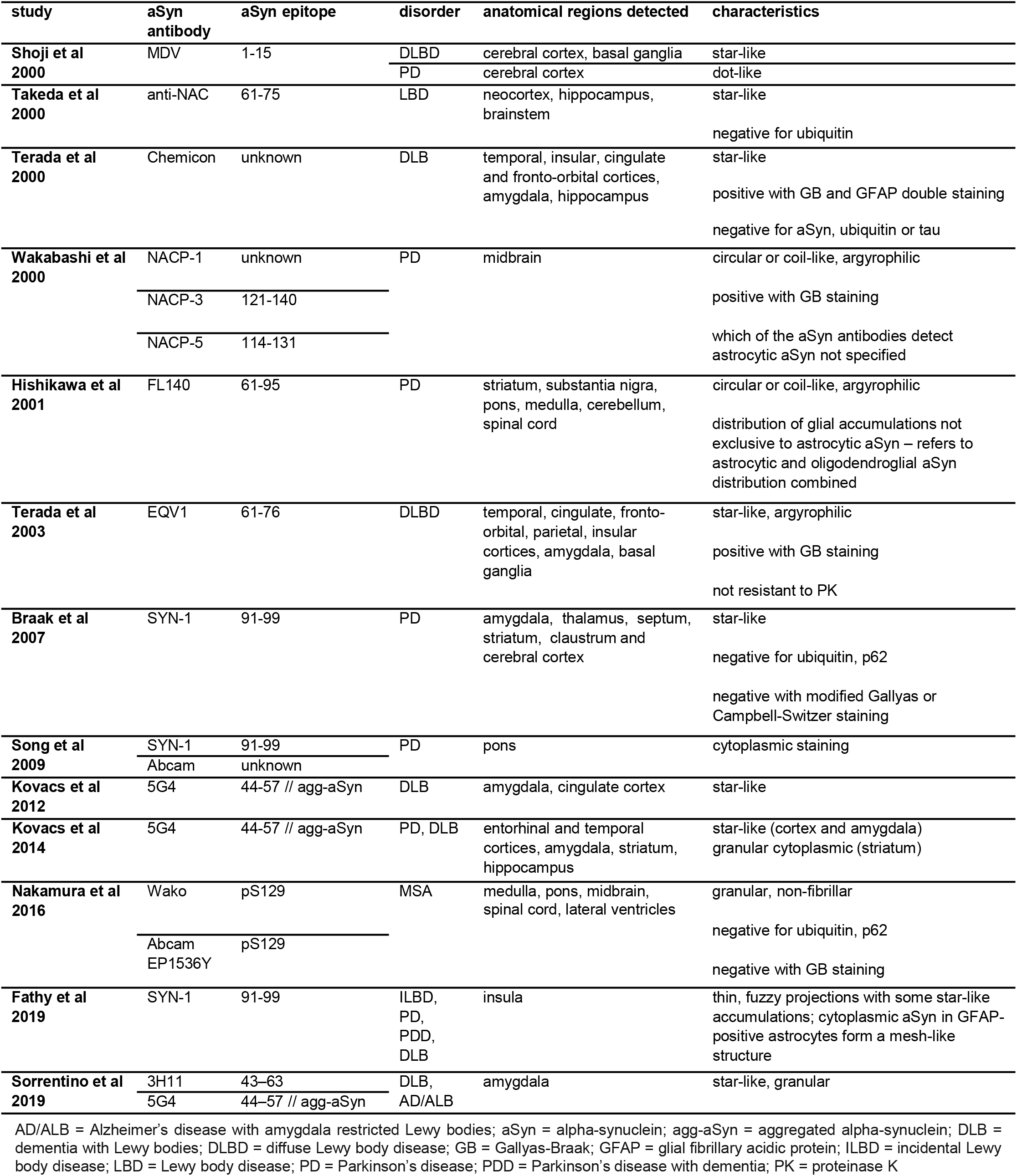
The studies since 2000 investigating the astrocytic aSyn pathology in human LB disease brains. AD/ALB = Alzheimer’s disease with amygdala restricted Lewy bodies; aSyn = alpha-synuclein; agg-aSyn = aggregated alpha-synuclein; DLB = dementia with Lewy bodies; DLBD = diffuse Lewy body disease; GB = Gallyas-Braak; GFAP = glial fibrillary acidic protein; ILBD = incidental Lewy body disease; LBD = Lewy body disease; PD = Parkinson’s disease; PDD = Parkinson’s disease with dementia; PK = proteinase K

One of the key reasons for the scarcity of data on astrocytic aSyn is that only a few of the aSyn antibodies appear to reveal these species. Although previous studies have reported that the astrocytic aSyn is detected by antibodies with epitopes against the NAC region of aSyn (Braak et al., 2007; Kovacs et al., 2012; Sorrentino et al., 2019), they did not define the sequence properties of aSyn or identify the molecular determinants underpinning their observations. These astrocytic aSyn species are not revealed by the classical inclusion markers such as positivity for ubiquitin (Braak et al., 2007; Kovacs et al., 2014; Nakamura et al., 2016; Takeda et al., 2000; Terada et al., 2000) and p62 (Braak et al., 2007; Kovacs et al., 2014; Nakamura et al., 2016; Takeda et al., 2000; Terada et al., 2000), and their post-translational modification (PTM) profile and aggregation states have not been systematically investigated beyond two studies that assessed S129 phosphorylation (pS129) status (Nakamura et al., 2016; Sorrentino et al., 2019). Furthermore, the antibodies used to characterise and diagnose aSyn pathology are often directed at the C-terminal domain of the protein, which could lead to the under-reporting of aSyn astrocytic pathology. The lack of appropriate tools and techniques that allow for the selective isolation and characterization of the astrocytic aSyn presents major challenges to defining their biochemical properties and relationship to other aSyn brain pathologies.

To address these challenges, we assessed aSyn astrocytic pathology using an expanded set of antibodies that target epitopes throughout the entire sequence of the protein and for the first time against all known aSyn PTMs (8 PTMs). To further characterize the aggregation state of aSyn species in the astroglia, we used antibodies against known markers of LBs (ubiquitin, p62), aSyn aggregate-specific antibodies, the amyloid-specific dye Amytracker, and investigated their resistance to proteolysis by proteinase K. With the aim of shedding light on the biochemical properties of astrocytic aSyn pathology across different regions and synucleinopathies, we extended our studies to several limbic brain regions, including the entorhinal cortex, hippocampus, cingulate cortex, insula and amygdala of PD, PDD and DLB; and the pons, putamen, cerebellum, frontal cortex and occipital cortex of MSA.

Our studies show that astrocytic aSyn accumulations occur extensively in the limbic regions of LB disease cases. These aSyn species are post-translationally modified at Tyrosine 39 (Y39) and cleaved at both the N- and C-termini of the protein, as evidenced by the fact that they are only detected with antibodies targeting epitopes approximately between residues 34-99. In addition to presenting new insight into the sequence, aggregation state and biochemical properties of astrocytic aSyn, our work provides a validated toolset that should enable a more systematic re-assessment of the role of astrocytic aSyn pathology in the development and progression of synucleinopathies. Our findings also emphasise the importance of using an appropriate and validated detection tools capable of capturing the diversity of aSyn species to map and characterize aSyn pathology in astrocytes and other cell types.

## RESULTS

### Mapping the astrocytic aSyn proteoform in LB disorders

We have previously shown that the use of antibodies against different regions and post-translational modifications of aSyn enables revealing the pathological diversity across synucleinopathies (Altay et al., 2022). Therefore, we sought to use an expanded antibody tool box to characterize the aSyn astrocytic pathology. A complete list of the antibodies used in this study and their epitopes is shown in Supplementary Table 1. DLB entorhinal cortex was stained using two antibodies against the N-terminal (epitopes 1-20 and 34-45), two against the NAC region (epitopes 80-96 and 91-99) and two against the C-terminal regions (epitopes 110-115 and 134-138) of aSyn (Figure 1A). Serial sections from the same region were also screened for aSyn post-translational modifications (PTMs), including Serine (at Ser87 and Ser129) and Tyrosine (at Tyr39, Tyr125, Tyr133 and Tyr136) phosphorylations, N-terminal nitration at Tyr39 (nY39) and C-terminal truncation at residue 120 (Figure 1B). Whilst all the antibodies against non-modified aSyn were able to detect LBs and LNs, only the N-terminal antibody LASH-BL 34-45 and the two NAC region antibodies LASH-BL 80-96 and BD SYN-1 (epitope 91-99) were able to reveal the star-like astroglial aSyn structures (Figure 1A). We confirmed the specificity of these three antibodies to aSyn by pre-adsorption treatment, after which the positivity to LBs and star-like structures was lost (Supplementary Figure 1). Strikingly, only the two antibodies against the aSyn PTMs in the N-terminal region of the protein, i.e. pY39 and nY39, but not the antibodies targeting the C-terminal aSyn PTMs, were positive for these astroglial structures (Figure 1B).

**Figure 1:**
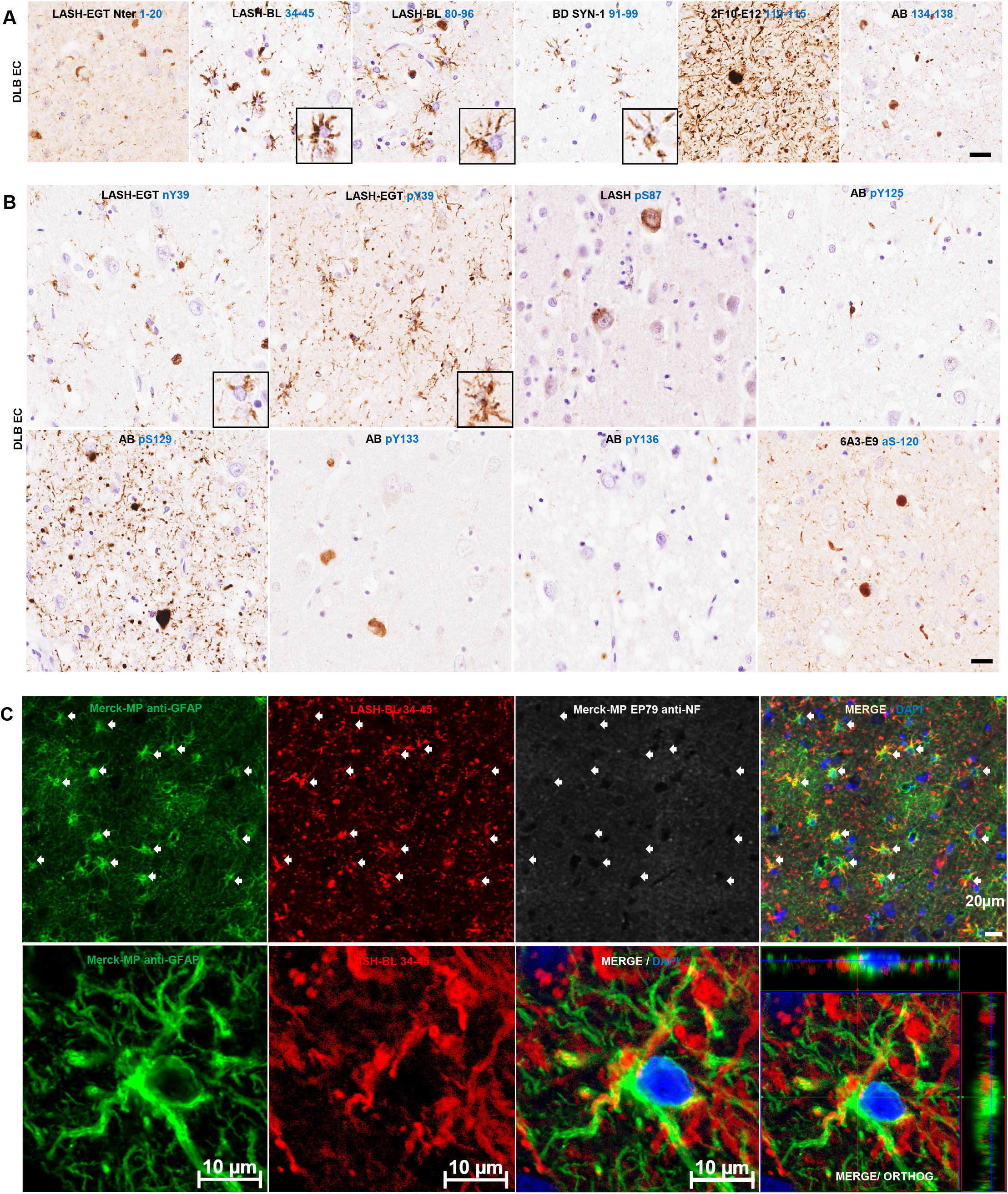
The star-like astrocytic aSyn accumulations in the entorhinal cortex (EC) of a DLB patient. (A) The EC of DLB brains were immunohistochemically stained using aSyn antibodies with epitopes against the N-terminus (LASH-EGTNter 1-20 and LASH-BL 34-45), the NAC region (LASH-BL 80-96 and BD SYN-1 91-99) and the C-terminus (2F10-E12 110-115 and AB 134-138) of aSyn. The extreme N-terminal antibody LASH-EGTNter as well as the C-terminal antibodies 2F10-E12 and AB 134-138 showed neuronal pathology in the soma and neurites. The late N-terminal antibody LASH-BL 34-45 as well as the two NAC region antibodies LASH-BL 80-96 and BD SYN-1 were positive for LBs and LNs, but also distinctively detected star-shaped glial aSyn species (insets). (B) The aSyn PTM antibodies against phosphorylation and nitration at Tyrosine 39 (Y39) were also reactive to the star-like astroglial pattern (insets). (C) Star-like aSyn species are associated with the GFAP-positive astrocytes in the DLB brains as shown by IF using antibodies for astrocytic and neuronal markers GFAP and NF, and LASH-BL 34-45 antibody against aSyn. The star-like aSyn species (arrows) appeared in and around the GFAP-positive astrocytes, and not in the LNs. Images on the upper panel taken using Olympus slide scanner at 40x magnification, and the lower panel on Zeiss LSM700 confocal microscope. Scale bar for Figure 1A-B is 20μm for the main images and 40μm for the insets. aSyn = alpha-synuclein; DLB = dementia with Lewy bodies; EC = entorhinal cortex; GFAP = glial fibrillary acidic protein; IF = immunofluorescence; LB = Lewy body; LN = Lewy neurite; NAC = non-amyloid component; NF = neurofilament; PTM = post-translational modification

To validate that these structures represent aSyn in the astrocytes, we performed immunofluorescent labelling. DLB cingulate cortex was stained using LASH-BL 34-45, the best performing antibody to reveal the star-like aSyn accumulations by immunohistochemistry (Figure 1A), and glial fibrillary acidic protein (GFAP) and neurofilament (NF) antibodies i.e. standard markers for astrocytes and neurons, respectively. The cortical LBs were positive for NF and LASH-BL 34-45, and negative for GFAP (Supplementary Figure 2A). The GFAP-positive astrocytes, on the other hand, were also positive for LASH-BL 34-45, and negative for NF by confocal imaging (Figure 1C). The oligodendroglia and the microglia in the white matter, marked with myelin basic protein (MBP) and ionised calcium binding adaptor protein 1 (Iba1) respectively, were negative to aSyn (Supplementary Figure 2B). Microglial positivity to aSyn was observed in the grey matter as rare events and did not show a star-like morphology (Supplementary Figure 2B). Collectively, these observations demonstrate that the star-like structures revealed by immunohistochemistry using the late N-terminal and NAC region aSyn antibodies represent aSyn compartmentalisation in astrocytes. These findings also suggest that the extreme N- and C-terminal regions of aSyn are either masked by heavy modifications, bound to other molecules or are cleaved, and thus explain why astrocytic aSyn cannot be detected with antibodies targeting these regions.

### Astrocytic aSyn is modified at Tyrosine 39

As shown in Figure 1B, our data show for the first time that astrocytic aSyn accumulations contain a mixture of aSyn species that are phosphorylated or nitrated at Tyrosine 39 (Y39). To corroborate our findings, we first validated the specificity of the aSyn pY39 and nY39 antibodies. The antibodies were incubated with aSyn recombinant protein bearing either pY39 or nY39 before using them to detect site-specifically nitrated and phosphorylated recombinant aSyn by slot blot (SB). The positive signal for aSyn nY39 and for aSyn pY39 recombinant proteins were lost after the pre-blocking of LASH-EGT nY39 and LASH-BL pY39 antibodies, respectively (Figure 2A). The same pre-blocking protocol was then repeated, and the pre-blocked antibody solutions applied onto DLB cingulate cortex tissues. Similarly, following the pre-blocking of the antibodies, the positivity for aSyn nY39 and pY39-positive species, including that for the astroglial structures, was abolished (Figure 2B). Next, we immunofluorescently co-labelled the cingulate sections with GFAP and aSyn pY39 or nY39 antibodies. Consistent with our observations by brightfield microscopy (Figure 1B), GFAP-positive astrocytes showed star-like aSyn inclusions positive for nY39 (Figure 2C).

**Figure 2:**
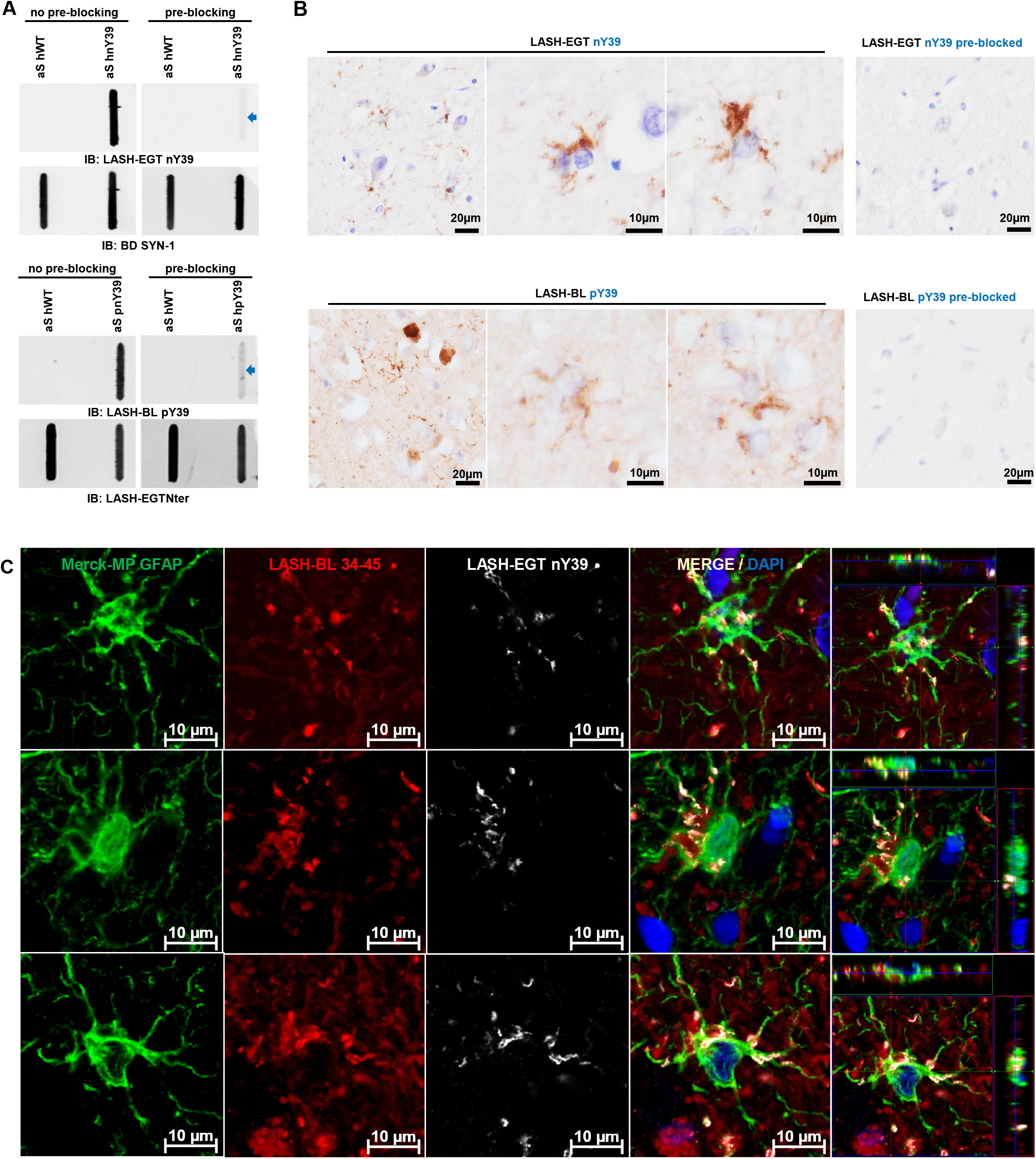
Astrocytic aSyn is modified at Tyrosine 39. (A) The specificity of the aSyn PTM antibodies against phosphorylation and nitration at Y39 were validated via antibody pre-blocking overnight followed by SB analysis. (B) The signal for astrocytic aSyn phosphorylated and nitrated at Y39 was lost with the antibody pre-blocking in the DLB cingulate cortex. (C) The GFAP- and aSyn-positive astrocytes in the cingulate cortex of DLB were positive for aSyn nY39. aSyn = alpha-synuclein; DLB = dementia with Lewy bodies; GFAP = glial fibrillary acidic protein; PTM = post-translational modification; SB = slot blot

### Astrocytic aSyn accumulations occur across LB disorders and may be truncated in the N- and C-termini

Having established that the star-like aSyn structures are astrocytic in DLB brains, we then sought to explore if the astrocytic aSyn exhibits similar biochemical and staining properties across other LB disorders. We screened, using the same set of antibodies against non-modified aSyn, the cingulate cortices of sporadic and familial PD, PDD and DLB cases. The astrocytic aSyn accumulations were observed widely across these LB diseases (Figure 3A; Supplementary Figure 3). These sections were double-labelled with aSyn LASH-BL 34-45 antibody and GFAP, which revealed that GFAP-positive astrocytes are positive to the aSyn accumulations in these cases (Supplementary Figure 4). The pons, putamen, cerebellum, frontal cortex and occipital cortex of MSA cases were stained using these aSyn antibodies, but we failed to detect any astroglial pathology in the MSA brains.

**Figure 3:**
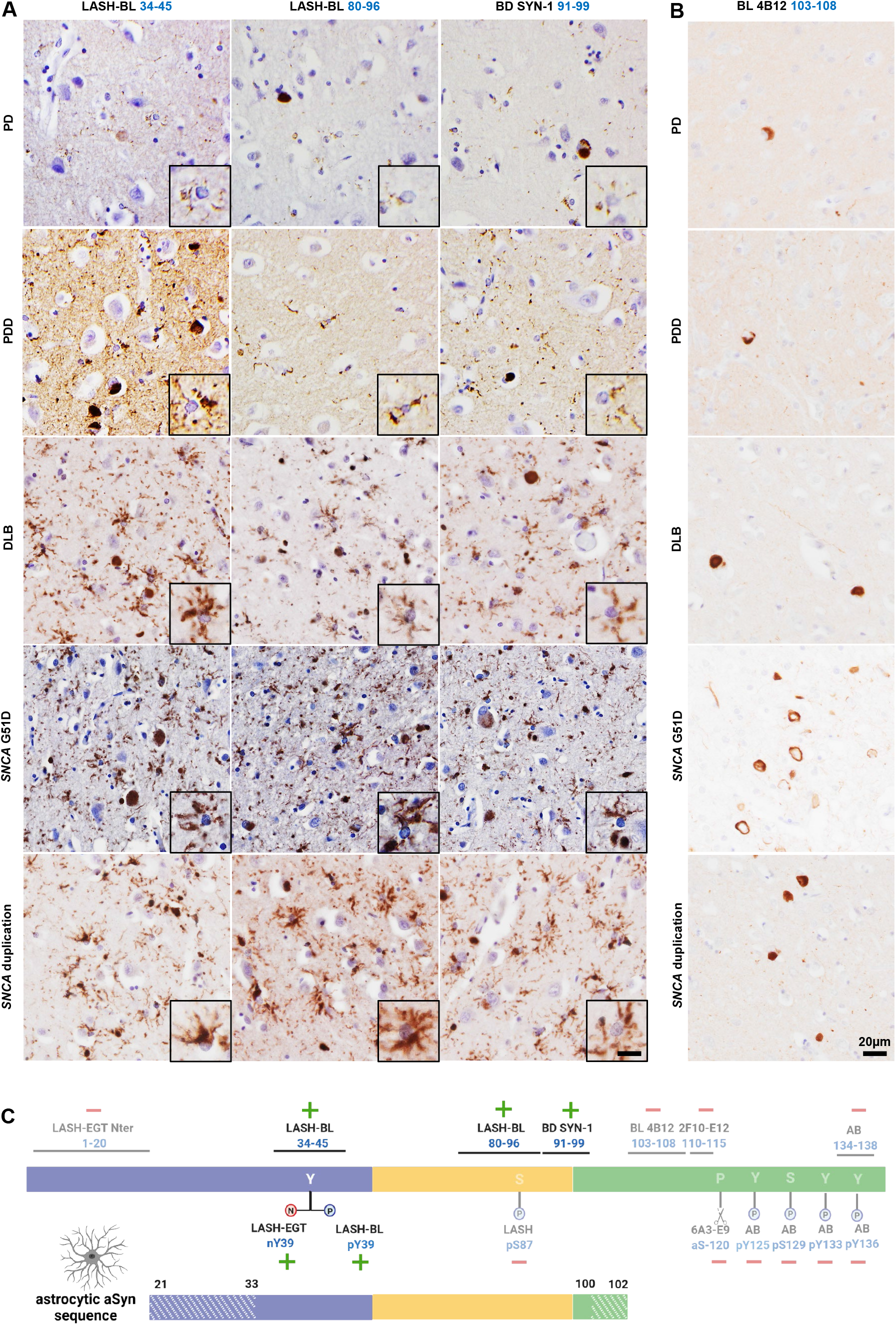
Astrocytic aSyn accumulations occur across LB disorders, and may be truncated in the N- and C-termini. (A) PD, PDD, DLB, *SNCA* G51D mutation and *SNCA* duplication cingulate cortices were immunohistochemically stained using three aSyn antibodies, LASH-BL 34-45, LASH-BL 80-96 and BD SYN-1, and astrocytic accumulations (insets) were revealed across these LB disorders. (B) To further map the C-terminal truncation region of the astrocytic aSyn, the same cingulate cortex sections were stained using the C-terminal BL 4B12 antibody with an epitope 103-108 of aSyn. Neuronal inclusions were revealed, but the astrocytic aSyn accumulation was not detected, suggesting that the aSyn species associated with the astrocytes are truncated at residues 21-33 in the N-terminus, and at residues 100-102 in the C-terminus. (C) A diagram to show the antibodies that are positive and negative for astrocytic aSyn, and their epitopes. The areas in stripes denote the potential truncation regions in the N- and C-termini. Schematic created with BioRender.com (agreement no: *DJ23GJF70T*). Scale bar for Figure 3A is 20μm for the main images and 40μm for the insets. aSyn = alpha-synuclein; DLB = dementia with Lewy bodies; LB = Lewy body; PD = Parkinson’s disease; PDD = Parkinson’s disease with dementia

To determine if the astrocytic aSyn species are also N-terminally and/or C-terminally truncated, and to more precisely map their sequence, we stained serial cingulate sections using BL 4B12 antibody, which targets residues 103-108. Interestingly, the astroglial structures were not detected using this antibody in any of the LB disorder cases in the cingulate cortex (Figure 3B). Our results suggest that astrocytic aSyn may be truncated in the N-terminus between residues 21-33, and in the C-terminus between residues 100-102 (Figure 3C).

### Astrocytic aSyn accumulations are not immunoreactive for the canonical aSyn aggregation markers

To investigate the nature and aggregation state of aSyn in these accumulations, the astrocytic aSyn species were screened for the classical markers of aSyn aggregation and inclusion formation. DLB cingulate cortex was triple labelled with LASH-BL 34-45 and GFAP, and either with antibodies against p62, ubiquitin, aSyn pS129 or with the amyloid dye Amytracker. In line with our brightfield microscopy results (Figure 1B), the cortical LBs and LNs showed strong positivity to aSyn pS129, whereas the astrocytes positive for LASH-BL 34-45 remained negative for aSyn pS129 (Figure 4A). Similarly, whereas the cortical LBs were positive for ubiquitin and p62, the aSyn-positive astrocytes were negative for these aggregate markers (Figure 4B-C). In contrast, astrocytic aSyn accumulations showed only partial positivity to amyloid dye Amytracker, which also labelled the cortical LBs (Figure 4D). Considering that several of the key lysine residues, found to be ubiquitinated in LBs (Anderson et al., 2006), reside in the N-terminal domain of the protein (i.e. K12, K21 and K23), and that p62 is a monoubiquitin- and polyubiquitin-binding protein (Cavey et al., 2004; Lee and Weihl, 2017; Raasi et al., 2005), the absence of ubiquitin and p62 positivity in the astrocytic aSyn is in line with our observations that the aSyn species in the astroglia are N-terminally truncated.

**Figure 4:**
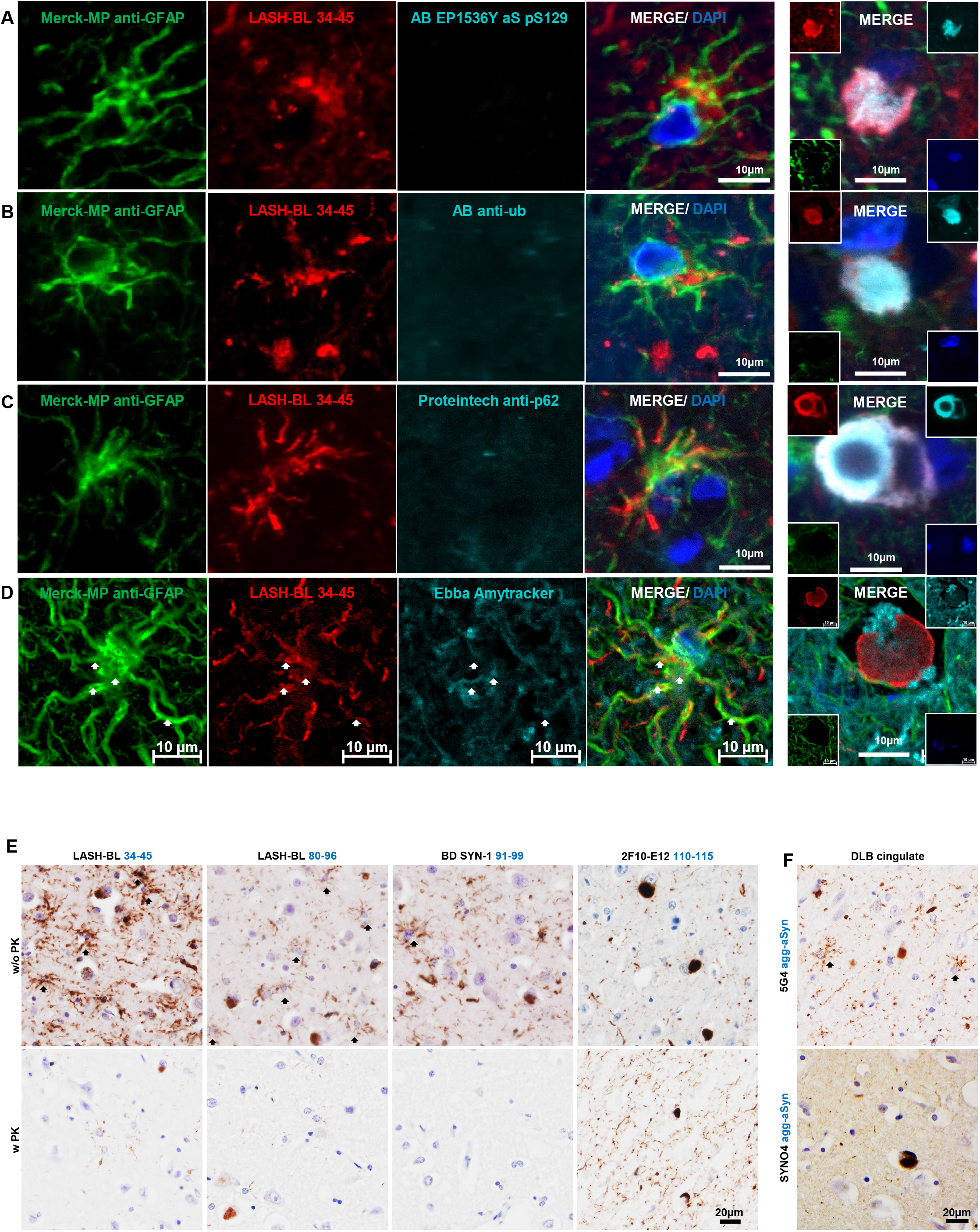
Astrocytic aSyn accumulations are free of the canonical aSyn aggregation markers. (A-C) The GFAP-positive astrocytic accumulations in DLB cingulate cortex were negative to aSyn pS129 (AB EP1536Y), ubiquitin and p62. (D) These astrocytic species showed partial overlap with the amyloid marker Amytracker (arrows). Images of cortical LBs are included as positive controls for aSyn pS129, ubiquitin, p62 and Amytracker reactivity. Images for Figure 4A-C taken using Olympus slide scanner at 40x magnification, and for Figure 4D on Zeiss LSM700 confocal microscope. (E) The astrocytic aSyn signal (arrows) was largely abolished after PK treatment in DLB cingulate cortex. The 2F10-E12 staining was included as a positive control to show the PK resistance of LBs and LNs. (F) The star-shaped astrocytic aSyn accumulations were revealed by the 5G4 antibody, but not by the SYNO4 antibody in the DLB cingulate cortex. agg-aSyn = aggregated alpha-synuclein; aSyn = alpha-synuclein; DLB = dementia with Lewy bodies; GFAP = glial fibrillary acidic protein; LB = Lewy body; LN = Lewy neurite; PK = proteinase K

One of the characteristics of aggregated aSyn in LB diseases is their resistance to proteinase K (PK) digestion (Neumann et al., 2004; Tanji et al., 2010). To further characterize the aggregation state of aSyn in astrocytes, we treated the DLB cingulate cortex tissues with PK, and observed that the large majority of the astrocytic aSyn signal disappeared after PK treatment (Figure 4E). Next, we profiled the astrocytic aSyn using two antibodies, 5G4 and SYNO4, that show preferential binding to aggregated aSyn (Kovacs et al., 2014, 2012; Kumar et al., 2020; Vaikath et al., 2015). Interestingly, the star-shaped astrocytic aSyn accumulations were revealed by the 5G4 antibody, but not by the SYNO4 antibody (Figure 4F). Altogether, these observations suggest that aSyn species that accumulate in the astrocytes may not possess the amyloid-like properties of aSyn fibrils found in LBs and LNs, but could still represent a mixture of soluble and non-amyloidogenic aggregates i.e. oligomers. Unfortunately, the lack of oligomer-specific antibodies and our inability to isolate and interrogate astrocytic aSyn make it difficult to more precisely determine the exact aggregation state of aSyn in the astrocytes.

### Astrocytic aSyn accumulations occur in several limbic regions of LB disorders

After having observed the frequent occurrence of astrocytic aSyn in the cingulate cortices of LB disorders (Figure 3A), we expanded our screening of astrocytic aSyn species in other limbic brain regions. We observed that the astrocytic aSyn accumulations were also prominently present in the entorhinal cortex, the insula, the amygdala and the hippocampus (Figure 5A; Supplementary Figure 5) of LB disorders. Interestingly, we identified different morphologies of astrocytic aSyn accumulations. The majority of the astrocytic aSyn accumulations appeared in soma-sparing star-like forms typically labelling the ramified processes of astrocytes (Figure 5B). In the hippocampal subregions, the astrocytic aSyn accumulations were predominantly in the soma and did not exhibit a star-shaped morphology (Figure 5C). Altogether, our findings demonstrate that astrocytic aSyn is a prominent pathological feature of LB diseases, and presents itself in a number of brain regions.

**Figure 5:**
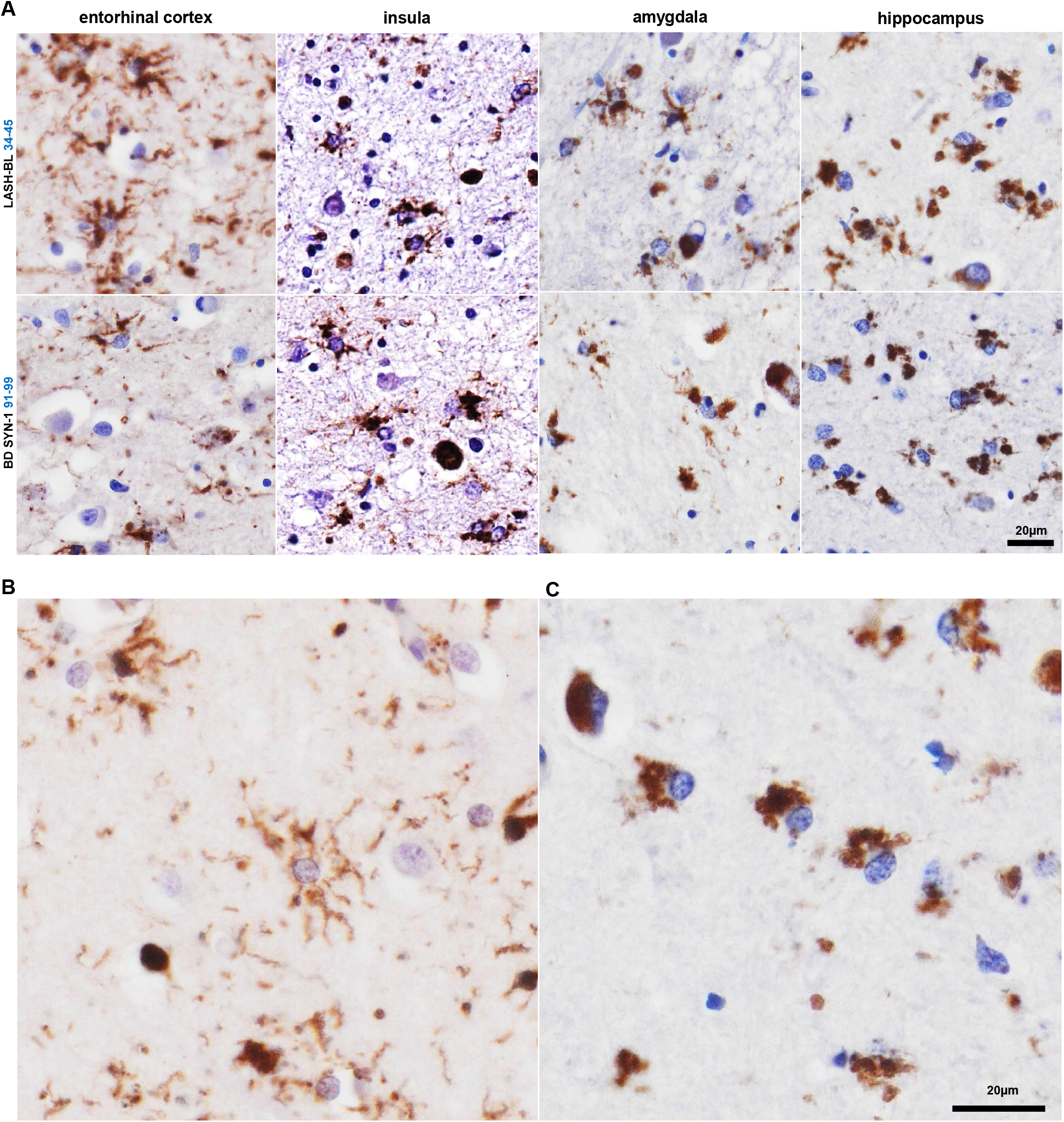
Astrocytic aSyn accumulations occur in several limbic regions of LB disease brains. (A) The astrocytic aSyn accumulations were encountered in the EC (*SNCA* G51D), insula (PDD), amygdala (DLB) and hippocampal CA4 (*SNCA* duplication) regions of LB disorders. (B) The astrocytic accumulations showed morphological diversity, with the majority showing a star shape and labelling the ramified processes (cingulate cortex). (C) Some of the astrocytic aSyn appeared as cytoplasmic accumulations (hippocampal CA4). Figure 5B-C images from a *SNCA* duplication brain stained with the LASH-BL 34-45 antibody. aSyn = alpha-synuclein; CA4 = cornu ammonis 4; DLB= dementia with Lewy bodies; EC = entorhinal cortex; LB = Lewy body; PD = Parkinson’s disease; PDD = Parkinson’s disease with dementia.

## DISCUSSION

Astrocytic aSyn pathology is a relatively less explored aspect of neuropathology in the synucleinopathies. In this study, we systematically characterised the biochemical properties and aggregation state of astrocytic aSyn accumulations using antibodies against non-modified and post-translationally modified forms of aSyn across the Lewy body diseases. Our results demonstrate that the astrocytic aSyn accumulations are widely present in several brain regions of PD, PDD and DLB cases, are negative for ubiquitin, p62 and aSyn pS129, but show positivity to other aSyn PTMs, including nitration and phosphorylation at Y39. Furthermore, only a subset of antibodies against non-modified aSyn are able to reveal astrocytic aSyn in brain tissues. These are antibodies that targeted the NAC (80-96 and 91-99) and the late N-terminal (34-45) regions of the protein. This is in line with the previous studies that reported astrocytic positivity only, or primarily with NAC region aSyn antibodies (Braak et al., 2007; Fathy et al., 2019; Hishikawa et al., 2001; Kovacs et al., 2014, 2012; Song et al., 2009; Sorrentino et al., 2019; Takeda et al., 2000; Terada et al., 2003) (Figure 6A). However, our studies allow for the more precise mapping, to the extent possible using antibodies, of the putative cleavage sites and demonstrate conclusively that the majority of astrocytic aSyn are both N- and C-terminally truncated (Figure 6B). In addition, we demonstrate for the first time that astrocytic aSyn exists as a mixture of non-amyloid species that are phosphorylated or nitrated at Y39.

**Figure 6:**
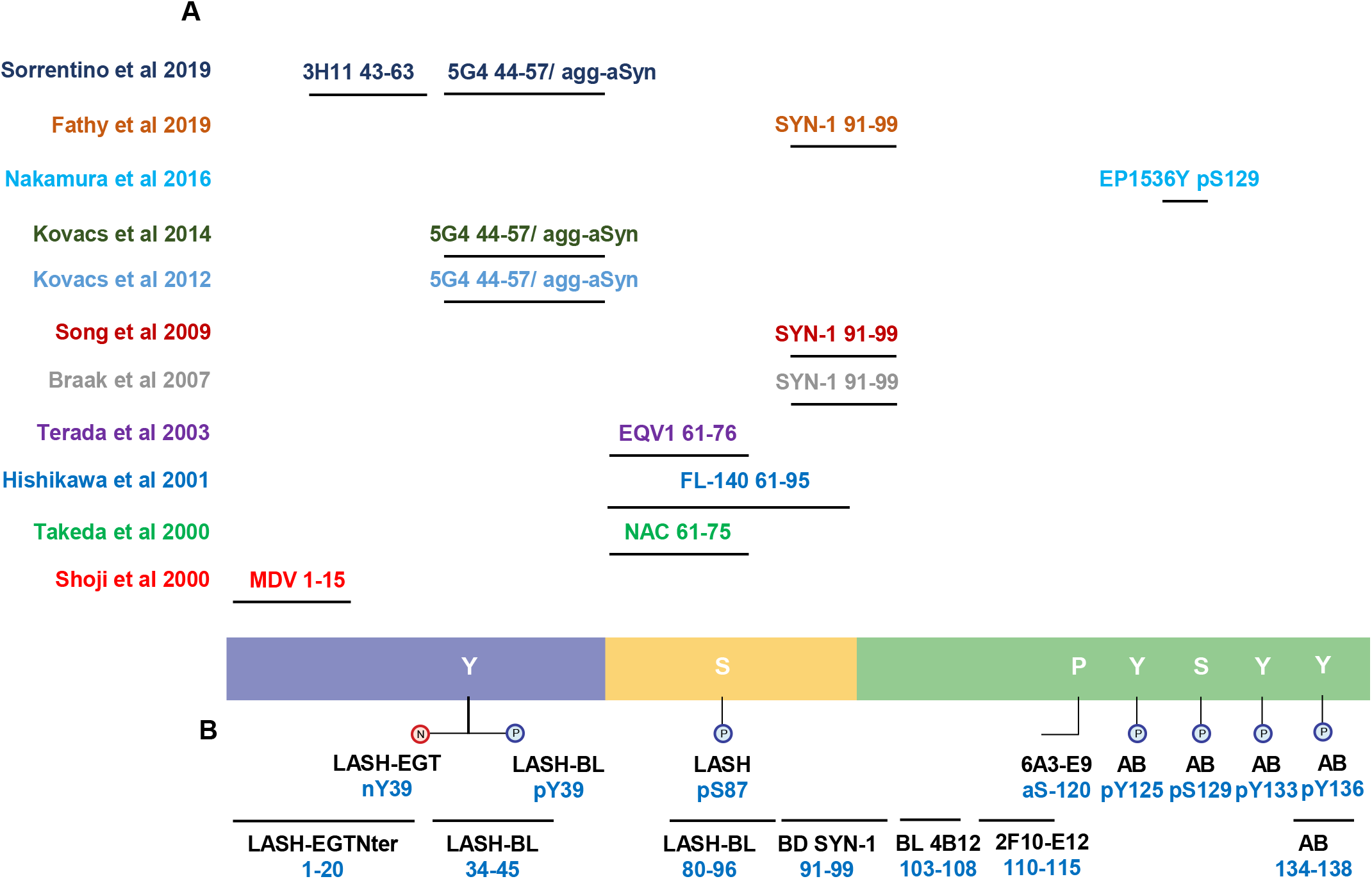
(A) A diagrammatic representation of the antibodies used in the previous publications reporting aSyn positivity in the astrocytes (only the studies using antibodies with a specified epitope are included). (B) The antibodies used in this study. aSyn = alpha-synuclein; agg-aSyn = aggregated alpha-synuclein

The fact that we were not able to detect astrocytic aSyn species with three antibodies targeting the C-terminal region of the protein spanning residues 103 to 138 strongly suggests that the astrocytic aSyn may be C-terminally truncated somewhere between residues 100-102. Similarly, the astrocytic aSyn species were detected by an antibody targeting residues 34-45 (LASH-BL 34-45), but not an antibody targeting the first N-terminal 20 amino acids (LASH-EGTNter 1-20). These results suggest that the N-terminal truncation of astrocytic aSyn is likely to occur between residues 21-33. Although an initial study (Shoji et al., 2000) reported the detection of astrocytic aSyn using an antibody with an N-terminal epitope (MDV, 1-15), subsequent studies showed that N-terminal antibodies covering aSyn residues 1-21 (Sorrentino et al., 2019; Takeda et al., 2000; Terada et al., 2003) did not detect astrocytic aSyn. Altogether, these data demonstrate that the great majority of aSyn in astrocytes are subjected to both N- and C-terminal cleavage at approximately residues 21-33 and 100-102, respectively. These results are supported by our findings that astrocytic aSyn inclusions were immunoreactive for only aSyn PTMs in the mid-N-terminal region. However, we cannot rule out the possibility that the extreme N- and C-terminal sequences are masked by aSyn interactions with other proteins, especially since both termini serve as hubs for regulating aSyn membranes/lipids and protein interactions. If this is the case, it would suggest that aSyn conformations and interactome in astrocytes are distinct from those in neurons, where aSyn is detectable using both N- and C-terminal-targeting antibodies. Whether truncated aSyn species are cleaved in the neurons, in the astrocytes or in the extracellular space is an important gap of knowledge that should be addressed and could shed new light into the role of PTMs in regulating the function/dysfunction of aSyn and mechanisms of aSyn trafficking between neurons and glia in the disease brains.

Very little is known about the aggregation state of the astrocytic aSyn species. Kovacs and colleagues (Kovacs et al., 2014, 2012) were the first to show positivity for astrocytes with the 5G4 antibody, reported to be specific for oligomeric and fibrillar forms of aSyn (Kovacs et al., 2012; Kumar et al., 2020). Similar astrocytic positivity using 5G4 was also detected by Sorrentino and colleagues (Sorrentino et al., 2019), but there has not been any studies to validate these findings using multiple aggregated aSyn antibodies and define the aggregation state of aSyn in astrocytes. Similarly, only two studies have assessed the ultrastructure of astrocytic aSyn by EM (Kovacs et al., 2014; Nakamura et al., 2016). Nakamura et al. described the aSyn pS129-positive subpial astrocytic processes in the MSA brains as non-filamentous (Nakamura et al., 2016), and Kovacs and colleagues reported that the astrocytic accumulations of LB diseases are beta-sheet-rich oligomers (Kovacs et al., 2014).

In this study, we investigated the aggregation state of aSyn in astrocytes using multiple approaches, including aSyn conformational/ aggregate-specific antibodies (5G4 and SYNO4), amyloid dyes and limited proteolysis (PK resistance). In line with Kovacs and colleagues (Kovacs et al., 2014), the astrocytic inclusions of LB diseases were revealed by 5G4, but were not detected with SYNO4. We also report that the majority of the astrocytic aSyn did not show resistance to PK digestion, and were only partially positive to the amyloid dye Amytracker. These findings, combined with our observation that astrocytic aSyn accumulations are not positive for the canonical markers of LBs, including ubiquitin, p62, and the most common pathology-associated aSyn PTM, pS129 (Anderson et al., 2006; Fujiwara et al., 2002), suggest that aSyn accumulations in astrocytes possess a distinct PTM and sequence signature, and may be composed primarily of oligomers or other non-fibrillar forms of the protein.

We speculate that the lack of aSyn phosphorylation at S129 is because the astrocytic aSyn species are truncated in the C-terminus, and no longer carry the binding site for the aSyn pS129 antibodies. Likewise, these aSyn accumulations are cleaved in the N-terminus, the domain where aSyn is found to be ubiquitinated in the disease brains (Anderson et al., 2006). Given that aSyn pS129 has been reported to be important for priming aSyn ubiquitination (Hasegawa et al., 2002), the absence of pS129 could explain the absence of ubiquitination at other lysine residues in the protein. One final possibility is that both of these aSyn PTMs are linked to the formation of aSyn pathology (Anderson et al., 2006; Fujiwara et al., 2002; Hasegawa et al., 2002), and their absence suggests that the astrocytic aSyn species exist in non-aggregated forms. The fact that astrocytic aSyn is cleaved and non-fibrillar at the same time is surprising given that removal of the solubilising N- and C-terminal domains is expected to increase the hydrophobicity and aggregation propensity of the protein (Bodles et al., 2000; Crowther et al., 1998; Eliezer et al., 2001; Giasson et al., 2001; Han et al., 1995; Volpicelli-Daley et al., 2011). Therefore, more extensive investigations of astrocytic aSyn conformations and aggregation state are needed. These studies could shed novel insights into the function(s) of aSyn in astrocytes and the role of astrocytic pathology in the pathogenesis of LB diseases. Furthermore, understanding what keeps these truncated aSyn species from forming fibrils in astrocytes could shed light on novel mechanisms for regulating aSyn aggregation.

These observations raise important questions about the origins and mechanisms involved in the astrocytic uptake, processing, degradation and/or release of aSyn. Cell culture studies have shown that astrocytes take up (Braidy et al., 2013; Cavaliere et al., 2017; Hua et al., 2019; H.-J. Lee et al., 2010b; Lindstrom et al., 2017; Loria et al., 2017; Rostami et al., 2017), degrade (Hua et al., 2019; Lindstrom et al., 2017; Loria et al., 2017; Rostami et al., 2017), and/or release (Cavaliere et al., 2017; Loria et al., 2017; Rostami et al., 2017) aSyn. Yet, a consensus has not been reached on whether or not the astrocytic uptake and processing of aSyn may have cytoprotective (Hua et al., 2019; Lindstrom et al., 2017; Loria et al., 2017) or cytotoxic (Braidy et al., 2013; Cavaliere et al., 2017; H.-J. Lee et al., 2010b; Lindstrom et al., 2017; Rostami et al., 2017) consequences. Kovacs and colleagues have shown that the astrocytic aSyn is localised in the endo-lysosomal compartments in the disease brains (Kovacs et al., 2014). Similarly, cell model-based research has shown that glial-glial and glial-neuronal oligomeric aSyn transfer can occur in lysosomal vesicles via direct transfer or tunnelling nanotubes (Cavaliere et al., 2017; Loria et al., 2017; Rostami et al., 2017). The fact that the great majority, if not all, of astrocytic aSyn across different LBDs is truncated suggests differential processing of aSyn in the astrocytes that may reflect its astrocytic functions, or a cellular response to aSyn species originating neurons or other glial cells. Altogether, a precise understanding of the astrocytic involvement in the cell-to-cell propagation of misfolded aSyn is needed to grasp the pathology spreading pathways in LB diseases.

Astrocytes are the most populous type of glial cells in the brain, with crucial functions in neuronal survival, synaptic maintenance, glucose metabolism, water homeostasis and in immune response (Sofroniew and Vinters, 2010). Insults may activate astrocytes (Wilhelmsson et al., 2006), which can in turn signal the microglia (Farina et al., 2007; H.-J. Lee et al., 2010a; Zhang et al., 2005) and act as key determinants of microglial activation and neuroinflammation in disease progression (Yamanaka et al., 2008). Furthermore, aSyn aggregates have been reported to activate both astrocytes (Chavarria et al., 2018; Chou et al., 2021; Fellner et al., 2013; Klegeris et al., 2006; H.-J. Lee et al., 2010b) and microglia (Fellner et al., 2013; E.-J. Lee et al., 2010) into giving an inflammatory response. Nitrated aSyn in particular has been reported to induce microglial activation (Reynolds et al., 2009, 2008a, 2008b; Thomas et al., 2007), which may then attain neurotoxic characteristics (Reynolds et al., 2009, 2008b). We found astrocytic aSyn to be nitrated at Y39, and speculate that this specific aSyn PTM may play a key role in the astrocytic signalling of microglia and neuroinflammation in LB diseases. Further studies to investigate the mechanisms of astrocytic activation of microglia, and the involvement of aSyn nY39 taken up and/or released by astrocytes within this context may further explain the interaction of neuroinflammation and neurodegeneration in LB disorders.

To our knowledge, this is the first study that examined the post-translational modifications profile (serine and tyrosine phosphorylations, tyrosine nitration and N- and C-terminal truncations) of astrocytic aSyn inclusions in Lewy body disorders. Although this hypothesis cannot be validated by biochemical profiling due to technical limitations to the isolation of astrocytic accumulations from the rest of the aSyn pathology, the failure of four different N- and C-terminal antibodies to detect astrocytic aSyn species strongly supports our conclusions on the biochemical properties of aSyn in astrocytic pathology. Furthermore, this is the first study reporting on aSyn brain pathological species that is composed primarily of truncated aSyn species. Previous studies have also shown high abundance of N- and C-terminally truncated aSyn species in the human brain (Anderson et al., 2006; Bhattacharjee et al., 2019; Kellie et al., 2015; Moors et al., 2021; Ohrfelt et al., 2011) and appendix (Killinger et al., 2018); however, in many of these studies the full-length protein remains highly abundant as the dominant species, as evidenced by the fact that antibodies against phosphorylated aSyn at S129 remain the primary tools used to monitor and quantify aSyn pathology in human brains and in animal models of synucleinopathies. The co-occurrence of N- and C-terminal truncation in the astrocytic aSyn without fibrillisation is particularly important, as the NAC region alone is known to be prone to aggregation by itself (Giasson et al., 2001). Which cell-specific mechanisms may prevent aSyn from forming aggregates in the astrocytes can have implications for understanding the cellular determinants of aSyn pathology formation and therapeutic applications.

Our findings raise several important questions that should be addressed in future studies to clarify 1) if these truncated species of aSyn become cleaved after internalisation by the astrocytes, or are internalised after being cleaved; 2) why these astrocytic aSyn inclusions appear in abundance in LBDs but are spared in MSA; 3) the precise nature of the aggregation state of aSyn in these astrocytes; and 4) the occurrence of astrocytic pathology in relation to LB disease staging, clinical progression and clinical phenotypes. The expanded toolset that we present here should facilitate these studies and advance our understanding of the function of astrocytic aSyn in health and disease.

## MATERIALS AND METHODS

### Antibodies

The primary and secondary antibodies used in this study are detailed in Supplementary Table 1.

### Human brain tissue samples

Post-mortem human brains stored at Queen Square Brain Bank (QSBB), Institute of Neurology in University College London (UCL), and Oxford Brain Bank (OBB), Nuffield Department of Clinical Neurosciences in University of Oxford, were collected in accordance with approved protocols by the London Multicentre Research Ethics Committee and the Ethics Committee of the University of Oxford (ref 15/SC/0639). All participants had given prior written informed consent for the brain donation. Both brain banks comply with the requirements of the Human Tissue Act 2004 and the Codes of Practice set by the Human Tissue Authority (HTA licence numbers 12198 for QSBB and 12217 for OBB). 3 cases of sporadic PD, MSA and familial PD with *SNCA* G51D mutation, 2 cases with PDD, 1 case with DLB and 1 case with PD with *SNCA* duplication were derived from QSBB, and 2 cases with sporadic PD, 2 cases with PDD and 2 cases with DLB from OBB were used in this study.

### Immunohistochemistry with 3,3’-diaminobenzidine (DAB) revelation and imaging

Formalin-fixed paraffin-embedded (FFPE) sections were dewaxed in xylene and rehydrated through decreasing concentrations of industrial denatured alcohol (IDA). Antigen retrieval was carried out for the appropriate antibody (Supplementary Table 1). Autoclaving (AC) was run at 121°C for 10 minutes in citrate buffer (pH6.0). For formic acid (FA) pre-treatment, tissues were incubated in 80-100% FA for 15 minutes (except for 5 minutes with 5G4) at room temperature (RT). For PK pre-treatment, tissues were incubated at 37°C for 5 minutes in 20μg/mL of PK diluted in TE-CaCl_2_ buffer (50mM Tris-base, 1mM EDTA, 5mM CaCl_2_, 0.5% Triton X-100, adjusted to pH8.0). Next, the sections were incubated for 30 minutes in 3% hydrogen peroxide in phosphate buffered saline (PBS) for quenching the endogenous peroxidase activity. Sections were briefly rinsed in distilled water and PBS, blocked in 10% foetal bovine serum (FBS) for 30 minutes at RT, and left at 4°C overnight for incubation with the primary antibodies. Subsequently, the sections were washed in PBS-Tween 0.1% (PBS-T; 3x 5 minutes) and incubated in the secondary antibody-horseradish peroxidase (HRP) complex as part of REAL EnVision detection system (Dako #K5007) for 1h at RT. Sections were rinsed in PBS-T (3x 5 minutes) before visualisation with 3,3’-diaminobenzidine (DAB), and counterstained with haematoxylin. Finally, they were dehydrated in increasing concentrations of IDA, cleared in xylene (3x 5 minutes) and mounted using distyrene plasticiser xylene (DPX).

### Immunofluorescent labelling and imaging

After the blocking in 10% FBS in PBS-T for 60 minutes at RT, sections were washed in PBS for 5 minutes and incubated for 1 minute in TrueBlack lipofuscin autofluorescence quencher (Biotium #23007) in 70% ethanol. The sections were washed in PBS (3x 5 minutes) and incubated in primary antibodies overnight at 4°C. They were rinsed in PBS (3x 5minutes) and incubated in secondary antibodies for 1h at RT in dark. Amytracker 680 (Ebba Biotech) was applied according to the manufacturer’s instructions, at a dilution of 1:1,000, and the tissue washed in PBS (3x 5 minutes). The slides were mounted using an aqueous mounting medium with DAPI (Vector Laboratories #H-1500-10). Tiled imaging was carried out on the Olympus VS120 microscope. Confocal imaging was carried out on a confocal laser-scanning microscope (LSM 700, Carl Zeiss, Germany), and image analysis on Zen Digital Imaging software (RRID: SCR_013672).

### Recombinant aSyn generation, antibody pre-adsorption and slot blot (SB) analysis

aSyn expression and purification was performed as described (Fauvet et al., 2012). In brief, aSyn human WT-encoding pT7-7 plasmids were used to transform BL21(DE3) chemically competent *E. coli*, which were then grown on an agar dish supplemented with ampicillin. A single colony was transferred to Luria broth (LB) media with ampicillin at 100μg/mL, the small culture was left to grow at 37 °C on shaker (at 180RPM) for 16h, and was then used to inoculate a large culture of 6L LB media supplemented with ampicillin at 100μg/mL. aSyn expression was induced at an optic density (OD_600_) of 0.5-0.6A, using isopropyl β-D-1-thiogalactopyranoside at a final concentration of 1mM. The culture was grown for another 4h on shaker, centrifuged at 4,000*g* for 15min at 4 °C, and the pellet collected. The lysis buffer of 20mM Tris pH8.0, 0.3μM phenylmethylsulfonyl fluoride (PMSF) protease inhibitor and cOmplete, mini, EDTA-free protease inhibitor cocktail tablet (Roche #4693159001; one tablet per 10mL lysis buffer) was used to re-suspend the pellet (10mL p/L of culture) on ice. Cell lysis was carried out by sonication (59s-pulse and 59s-no pulse over 5min at 60% amplitude), and the lysate was spun down for 30min at 20,000*g* and 4 °C. The supernatant was collected and boiled for 15min at 100 °C, and the centrifugation repeated before the supernatant was filtered via a 0.22μm syringe filter. The purification was performed by anion exchange chromatography and reverse-phase high performance liquid chromatography (HPLC). The protein quality control was carried out by liquid chromatography-mass spectrometry (LC-MS), ultra-performance liquid chromatography (UPLC), and SDS-PAGE separation and Coomassie staining. The preparation of the aSyn nY39 and pY39 proteins involved the use of a semisynthetic approach as described previously (Hejjaoui et al., 2012).

For the antibody pre-adsorption, 5-fold of recombinant aSyn protein, or just PBS as control, was added to the IHC-optimised antibody solution in PBS (see Supplementary Table 1 for the IHC dilutions). The mixture was incubated overnight at 4 °C on a wheel, and the probing protocol, adapted from (Kumar et al., 2020), was carried out for the slot blot analysis. 200ng of aSyn proteins diluted in PBS to 100μL were blotted on 0.22μm nitrocellulose membranes, which were blocked at 4 °C overnight in Odyssey blocking buffer (Li-Cor). After the incubation with primary antibodies diluted in PBS for 2h at RT, the membranes were washed x3 for 10min in PBS with 0.01% Tween-20 (PBS-T), incubated with the secondary antibodies diluted in PBS in the dark, and washed x3 for 10min in PBS-T. For the SB dilutions of the primary and secondary antibodies, see Supplementary Table 1. Imaging was carried out at 700nm and 800nm using Li-Cor Odyssey CLx, and the image processing using Image Studio 5.2.

---

**Supplementary Figure 1:** Specificity validation of the aSyn antibodies LASH-BL 34-45, LASH-BL 80-96 and BD SYN-1 by pre-adsorption followed by IHC on DLB cingulate cortex. aSyn = alpha-synuclein; DLB = dementia with Lewy bodies; IHC = immunohistochemistry

**Supplementary Figure 2:** (A) A representative image of a cortical LB positive for NF and LASH-BL 34-45, and negative for GFAP. Image from DLB cingulate cortex, taken using Zeiss LSM700 confocal microscope. (B) The oligodendrocytes and microglial cells, marked by anti-MBP and anti-Iba1 antibodies, respectively, were negative for aSyn in the white matter (upper two panels). Punctate aSyn positivity was detected in the microglial cells in the grey matter (lower panel) as rare events. Images taken from DLB cingulate cortex using Olympus slide scanner at 40x magnification. aSyn = alpha-synuclein; DLB = dementia with Lewy body; GFAP = glial fibrillary acidic protein; Iba1 = ionised calcium binding adaptor protein 1; MBP = myelin basic protein; LB = Lewy body; NF = neurofilament

**Supplementary Figure 3:** The cingulate cortex of sporadic PD, PDD, DLB, *SNCA* G51D mutation and *SNCA* duplication cases immunostained using antibodies against the N-terminal (LASH-EGTNter) and C-terminal (2F10-E12 and AB 134-138) of aSyn. Astrocytic aSyn was not detected using these antibodies. aSyn = alpha-synuclein; DLB = dementia with Lewy bodies; PD = Parkinson’s disease; PDD = Parkinson’s disease with dementia

**Supplementary Figure 4:** Representative IF images of GFAP-positive astrocytes from DLB, *SNCA G51D* mutation and *SNCA* duplication cingulate cortices, showing positivity for aSyn detected using LASH-BL 34-45 antibody. Images taken using Olympus slide scanner at 40x magnification. aSyn = alpha-synuclein; DLB = dementia with Lewy body; GFAP = glial fibrillary acidic protein; IF = immunofluorescence; NF = neurofilament

**Supplementary Figure 5:** Representative images from the *SNCA* duplication EC, insula and amygdala immunostained using antibodies against the N-terminal, NAC and C-terminal regions of aSyn. The astrocytic aSyn detected only by LASH-BL34-45, LASH-BL80-96 and BD SYN-1 (91-99) antibodies. Similar staining patterns were observed in the same regions from PD, PDD, DLB and *SNCA* G51D cases stained with the same antibody set. aSyn = alpha-synuclein; DLB = dementia with Lewy bodies; EC = entorhinal cortex; PD = Parkinson’s disease; PDD = Parkinson’s disease with dementia

**Supplementary Table 1:** The primary and secondary antibodies included in this study. AC = autoclave; agg-aSyn = aggregated alpha-synuclein; aSyn = alpha-synuclein; FA = formic acid; GFAP = glial fibrillary acidic protein; Iba1 = ionised calcium binding adaptor protein 1; IF = immunofluorescence; IHC = immunohistochemistry; MBP = myelin basic protein; mc = monoclonal; mus = mouse; na = not applicable; NF = neurofilament; pc = polyclonal; rab = rabbit; SB = slot blot

## Contributions of the authors

HAL and LP conceived and conceptualised the study. HAL, LP and MFA designed the experiments. JLH selected the cases originating the Queen Square Brain Bank, and LP those originating the Oxford Brain Bank. JLH contributed to, and LP and MFA finalised the determination of optimal IHC conditions for the aSyn antibodies. LP and AKLL contributed to the analysis and interpretation of neuropathological data. MFA performed all other experiments. MFA wrote the first draft of the manuscript, which was then reviewed and modified by all authors.

AKLL: Alan King Lun Liu

HAL: Hilal A. Lashuel

JLH: Janice L. Holton

LP: Laura Parkkinen

MFA: Melek Firat Altay

## Acknowledgments

This work was support by grants from the Swiss National Science Foundation (31ER30_186198), Michael J Fox Foundation (MJFF-020698) and EPFL. The Queen Square Brain Bank is supported by the Reta Lila Weston Institute of Neurological Studies, UCL Queen Square Institute of Neurology. The Oxford Brain Bank is supported by the Medical Research Council (MRC), Brains for Dementia Research (BDR) (Alzheimer Society and Alzheimer Research UK), Autistica UK and the NIHR Oxford Biomedical Research Centre. We thank Catherine Strand (QSBB, UCL) for the preparation of the requested sections, and Selene Lee and Livia Civitelli (Oxford) for their contributions to the immunohistochemical data collection. We thank all the donors and their families for the invaluable brain donation to the brain banks.

## REFERENCES

Altay, M.F., Kumar, S.T., Burtscher, J., Jagannath, S., Strand, C., Miki, Y., Parkkinen, L., Holton, J.L., Lashuel, H.A., 2022. Development and validation of an expanded antibody toolset that captures alpha-synuclein pathological diversity in Lewy body diseases. bioRxiv. https://doi.org/10.1101/2022.05.26.493598

Anderson, J.P., Walker, D.E., Goldstein, J.M., de Laat, R., Banducci, K., Caccavello, R.J., Barbour, R., Huang, J., Kling, K., Lee, M., Diep, L., Keim, P.S., Shen, X., Chataway, T., Schlossmacher, M.G., Seubert, P., Schenk, D., Sinha, S., Gai, W.P., Chilcote, T.J., 2006. Phosphorylation of Ser-129 Is the Dominant Pathological Modification of α-Synuclein in Familial and Sporadic Lewy Body Disease. J. Biol. Chem. 281, 29739–29752. https://doi.org/10.1074/jbc.M600933200

Baba, M., Nakajo, S., Tu, P.H., Tomita, T., Nakaya, K., Lee, V.M., Trojanowski, J.Q., Iwatsubo, T., 1998. Aggregation of alpha-synuclein in Lewy bodies of sporadic Parkinson’s disease and dementia with Lewy bodies. Am J Pathol 152, 879–884.

Bhattacharjee, P., Öhrfelt, A., Lashley, T., Blennow, K., Brinkmalm, A., Zetterberg, H., 2019. Mass Spectrometric Analysis of Lewy Body-Enriched α-Synuclein in Parkinson’s Disease. J. Proteome Res. 18, 2109–2120. https://doi.org/10.1021/acs.jproteome.8b00982

Bodles, A.M., Guthrie, D.J.S., Harriott, P., Campbell, P., Irvine, G.B., 2000. Toxicity of non-Aβ component of Alzheimer’s disease amyloid, and N-terminal fragments thereof, correlates to formation of β-sheet structure and fibrils: Toxicity of non-Aβ component and fragments thereof. European Journal of Biochemistry 267, 2186–2194. https://doi.org/10.1046/j.1432-1327.2000.01219.x

Braak, H., Sastre, M., Del Tredici, K., 2007. Development of α-synuclein immunoreactive astrocytes in the forebrain parallels stages of intraneuronal pathology in sporadic Parkinson’s disease. Acta Neuropathol 114, 231–241. https://doi.org/10.1007/s00401-007-0244-3

Braidy, N., Gai, W.-P., Xu, Y.H., Sachdev, P., Guillemin, G.J., Jiang, X.-M., Ballard, J.W.O., Horan, M.P., Fang, Z.M., Chong, B.H., Chan, D.Y., 2013. Uptake and mitochondrial dysfunction of alpha-synuclein in human astrocytes, cortical neurons and fibroblasts. Translational Neurodegeneration 2, 20. https://doi.org/10.1186/2047-9158-2-20

Cavaliere, F., Cerf, L., Dehay, B., Ramos-Gonzalez, P., De Giorgi, F., Bourdenx, M., Bessede, A., Obeso, J.A., Matute, C., Ichas, F., Bezard, E., 2017. In vitro α-synuclein neurotoxicity and spreading among neurons and astrocytes using Lewy body extracts from Parkinson disease brains. Neurobiology of Disease 103, 101–112. https://doi.org/10.1016/j.nbd.2017.04.011

Cavey, J.R., Ralston, S.H., Hocking, L.J., Sheppard, P.W., Ciani, B., Searle, M.S., Layfield, R., 2004. Loss of Ubiquitin-Binding Associated With Paget’s Disease of Bone p62 (SQSTM1) Mutations. J Bone Miner Res 20, 619–624. https://doi.org/10.1359/JBMR.041205

Chavarria, C., Rodríguez-Bottero, S., Quijano, C., Cassina, P., Souza, J.M., 2018. Impact of monomeric, oligomeric and fibrillar alpha-synuclein on astrocyte reactivity and toxicity to neurons. Biochemical Journal 475, 3153–3169. https://doi.org/10.1042/BCJ20180297

Chou, T.-W., Chang, N.P., Krishnagiri, M., Patel, A.P., Lindman, M., Angel, J.P., Kung, P.-L., Atkins, C., Daniels, B.P., 2021. Fibrillar α-synuclein induces neurotoxic astrocyte activation via RIP kinase signaling and NF-κB. Cell Death Dis 12, 756. https://doi.org/10.1038/s41419-021-04049-0

Crowther, R.A., Jakes, R., Spillantini, M.G., Goedert, M., 1998. Synthetic filaments assembled from C-terminally truncated α-synuclein. FEBS Letters 436, 309–312. https://doi.org/10.1016/S0014-5793(98)01146-6

Eliezer, D., Kutluay, E., Bussell, R., Browne, G., 2001. Conformational properties of α-synuclein in its free and lipid-associated states 1 1Edited by P. E. Wright. Journal of Molecular Biology 307, 1061–1073. https://doi.org/10.1006/jmbi.2001.4538

Farina, C., Aloisi, F., Meinl, E., 2007. Astrocytes are active players in cerebral innate immunity. Trends in Immunology 28, 138–145. https://doi.org/10.1016/j.it.2007.01.005

Fathy, Y.Y., Jonker, A.J., Oudejans, E., Jong, F.J.J., Dam, A. -M. W., Rozemuller, A.J.M., Berg, W.D.J., 2019. Differential insular cortex subregional vulnerability to α-synuclein pathology in Parkinson’s disease and dementia with Lewy bodies. Neuropathol Appl Neurobiol 45, 262–277. https://doi.org/10.1111/nan.12501

Fauvet, B., Mbefo, M.K., Fares, M.-B., Desobry, C., Michael, S., Ardah, M.T., Tsika, E., Coune, P., Prudent, M., Lion, N., Eliezer, D., Moore, D.J., Schneider, B., Aebischer, P., El-Agnaf, O.M., Masliah, E., Lashuel, H.A., 2012. α-Synuclein in Central Nervous System and from Erythrocytes, Mammalian Cells, and *Escherichia coli* Exists Predominantly as Disordered Monomer. J. Biol. Chem. 287, 15345–15364. https://doi.org/10.1074/jbc.M111.318949

Fellner, L., Irschick, R., Schanda, K., Reindl, M., Klimaschewski, L., Poewe, W., Wenning, G.K., Stefanova, N., 2013. Toll-like receptor 4 is required for α-synuclein dependent activation of microglia and astroglia. Glia 61, 349–360. https://doi.org/10.1002/glia.22437

Fujiwara, H., Hasegawa, M., Dohmae, N., Kawashima, A., Masliah, E., Goldberg, M.S., Shen, J., Takio, K., Iwatsubo, T., 2002. α-Synuclein is phosphorylated in synucleinopathy lesions. Nat Cell Biol 4, 160–164. https://doi.org/10.1038/ncb748

Giasson, B.I., Murray, I.V.J., Trojanowski, J.Q., Lee, V.M.-Y., 2001. A Hydrophobic Stretch of 12 Amino Acid Residues in the Middle of α-Synuclein Is Essential for Filament Assembly. J. Biol. Chem. 276, 2380–2386. https://doi.org/10.1074/jbc.M008919200

Han, H., Weinreb, P.H., Lansbury, P.T., 1995. The core Alzheimer’s peptide NAC forms amyloid fibrils which seed and are seeded by β-amyloid: is NAC a common trigger or target in neurodegenerative disease? Chemistry & Biology 2, 163–169. https://doi.org/10.1016/1074-5521(95)90071-3

Hasegawa, M., Fujiwara, H., Nonaka, T., Wakabayashi, K., Takahashi, H., Lee, V.M.-Y., Trojanowski, J.Q., Mann, D., Iwatsubo, T., 2002. Phosphorylated α-Synuclein Is Ubiquitinated in α-Synucleinopathy Lesions. J. Biol. Chem. 277, 49071–49076. https://doi.org/10.1074/jbc.M208046200

Hejjaoui, M., Butterfield, S., Fauvet, B., Vercruysse, F., Cui, J., Dikiy, I., Prudent, M., Olschewski, D., Zhang, Y., Eliezer, D., Lashuel, H.A., 2012. Elucidating the Role of C-Terminal Post-Translational Modifications Using Protein Semisynthesis Strategies: α-Synuclein Phosphorylation at Tyrosine 125. J. Am. Chem. Soc. 134, 5196–5210. https://doi.org/10.1021/ja210866j

Hishikawa, N., Hashizume, Y., Yoshida, M., Sobue, G., 2001. Widespread occurrence of argyrophilic glial inclusions in Parkinson’s disease. Neuropathol Appl Neurobiol 27, 362–372. https://doi.org/10.1046/j.1365-2990.2001.00345.x

Hua, J., Yin, N., Xu, S., Chen, Q., Tao, T., Zhang, J., Ding, J., Fan, Y., Hu, G., 2019. Enhancing the Astrocytic Clearance of Extracellular α-Synuclein Aggregates by Ginkgolides Attenuates Neural Cell Injury. Cell Mol Neurobiol 39, 1017–1028. https://doi.org/10.1007/s10571-019-00696-2

Kellie, J.F., Higgs, R.E., Ryder, J.W., Major, A., Beach, T.G., Adler, C.H., Merchant, K., Knierman, M.D., 2015. Quantitative Measurement of Intact Alpha-Synuclein Proteoforms from Post-Mortem Control and Parkinson’s Disease Brain Tissue by Intact Protein Mass Spectrometry. Sci Rep 4, 5797. https://doi.org/10.1038/srep05797

Killinger, B.A., Madaj, Z., Sikora, J.W., Rey, N., Haas, A.J., Vepa, Y., Lindqvist, D., Chen, H., Thomas, P.M., Brundin, P., Brundin, L., Labrie, V., 2018. The vermiform appendix impacts the risk of developing Parkinson’s disease. Science Translational Medicine 10, eaar5280. https://doi.org/10.1126/scitranslmed.aar5280

Klegeris, A., Giasson, B.I., Zhang, H., Maguire, J., Pelech, S., McGeer, P.L., 2006. Alpha-synuclein and its disease-causing mutants induce ICAM-1 and IL-6 in human astrocytes and astrocytoma cells. FASEB j. 20, 2000–2008. https://doi.org/10.1096/fj.06-6183com

Kovacs, G.G., Breydo, L., Green, R., Kis, V., Puska, G., Lőrincz, P., Perju-Dumbrava, L., Giera, R., Pirker, W., Lutz, M., Lachmann, I., Budka, H., Uversky, V.N., Molnár, K., László, L., 2014. Intracellular processing of disease-associated α-synuclein in the human brain suggests prion-like cell-to-cell spread. Neurobiology of Disease 69, 76–92. https://doi.org/10.1016/j.nbd.2014.05.020

Kovacs, G.G., Wagner, U., Dumont, B., Pikkarainen, M., Osman, A.A., Streichenberger, N., Leisser, I., Verchère, J., Baron, T., Alafuzoff, I., Budka, H., Perret-Liaudet, A., Lachmann, I., 2012. An antibody with high reactivity for disease-associated α-synuclein reveals extensive brain pathology. Acta Neuropathol 124, 37–50. https://doi.org/10.1007/s00401-012-0964-x

Kumar, S.T., Jagannath, S., Francois, C., Vanderstichele, H., Stoops, E., Lashuel, H.A., 2020. How specific are the conformation-specific α-synuclein antibodies? Characterization and validation of 16 α-synuclein conformation-specific antibodies using well-characterized preparations of α-synuclein monomers, fibrils and oligomers with distinct structures and morphology. Neurobiology of Disease 146, 105086. https://doi.org/10.1016/j.nbd.2020.105086

Lee, E.-J., Woo, M.-S., Moon, P.-G., Baek, M.-C., Choi, I.-Y., Kim, W.-K., Junn, E., Kim, H.-S., 2010. α-Synuclein Activates Microglia by Inducing the Expressions of Matrix Metalloproteinases and the Subsequent Activation of Protease-Activated Receptor-1. J.I. 185, 615–623. https://doi.org/10.4049/jimmunol.0903480

Lee, H.-J., Kim, C., Lee, S.-J., 2010a. Alpha-Synuclein Stimulation of Astrocytes: Potential Role for Neuroinflammation and Neuroprotection. Oxidative Medicine and Cellular Longevity 3, 283–287. https://doi.org/10.4161/oxim.3.4.12809

Lee, H.-J., Suk, J.-E., Patrick, C., Bae, E.-J., Cho, J.-H., Rho, S., Hwang, D., Masliah, E., Lee, S.-J., 2010b. Direct Transfer of α-Synuclein from Neuron to Astroglia Causes Inflammatory Responses in Synucleinopathies*. Journal of Biological Chemistry 285, 9262–9272. https://doi.org/10.1074/jbc.M109.081125

Lee, Y., Weihl, C.C., 2017. Regulation of SQSTM1/p62 via UBA domain ubiquitination and its role in disease. Autophagy 13, 1615–1616. https://doi.org/10.1080/15548627.2017.1339845

Lindstrom, V., Gustafsson, G., Sanders, L.H., Howlett, E.H., Sigvardson, J., Kasrayan, A., Ingelsson, M., Bergström, J., Erlandsson, A., 2017. Extensive uptake of α-synuclein oligomers in astrocytes results in sustained intracellular deposits and mitochondrial damage. Molecular and Cellular Neuroscience 82, 143–156. https://doi.org/10.1016/j.mcn.2017.04.009

Loria, F., Vargas, J.Y., Bousset, L., Syan, S., Salles, A., Melki, R., Zurzolo, C., 2017. α-Synuclein transfer between neurons and astrocytes indicates that astrocytes play a role in degradation rather than in spreading. Acta Neuropathol 134, 789–808. https://doi.org/10.1007/s00401-017-1746-2

Moors, T.E., Maat, C.A., Niedieker, D., Mona, D., Petersen, D., Timmermans-Huisman, E., Kole, J., El-Mashtoly, S.F., Spycher, L., Zago, W., Barbour, R., Mundigl, O., Kaluza, K., Huber, S., Hug, M.N., Kremer, T., Ritter, M., Dziadek, S., Geurts, J.J.G., Gerwert, K., Britschgi, M., van de Berg, W.D.J., 2021. The subcellular arrangement of alpha-synuclein proteoforms in the Parkinson’s disease brain as revealed by multicolor STED microscopy. Acta Neuropathol. https://doi.org/10.1007/s00401-021-02329-9

Nakamura, K., Mori, F., Kon, T., Tanji, K., Miki, Y., Tomiyama, M., Kurotaki, H., Toyoshima, Y., Kakita, A., Takahashi, H., Yamada, M., Wakabayashi, K., 2016. Accumulation of phosphorylated α-synuclein in subpial and periventricular astrocytes in multiple system atrophy of long duration: Phosphorylated α-synuclein in MSA astrocytes. Neuropathology 36, 157–167. https://doi.org/10.1111/neup.12243

Neumann, M., Müller, V., Kretzschmar, H.A., Haass, C., Kahle, P.J., 2004. Regional Distribution of Proteinase K-Resistant α-Synuclein Correlates with Lewy Body Disease Stage. J Neuropathol Exp Neurol 63, 1225–1235. https://doi.org/10.1093/jnen/63.12.1225

Ohrfelt, A., Zetterberg, H., Andersson, K., Persson, R., Secic, D., Brinkmalm, G., Wallin, A., Mulugeta, E., Francis, P.T., Vanmechelen, E., Aarsland, D., Ballard, C., Blennow, K., Westman-Brinkmalm, A., 2011. Identification of Novel α-Synuclein Isoforms in Human Brain Tissue by using an Online NanoLC-ESI-FTICR-MS Method. Neurochem Res 36, 2029–2042. https://doi.org/10.1007/s11064-011-0527-x

Raasi, S., Varadan, R., Fushman, D., Pickart, C.M., 2005. Diverse polyubiquitin interaction properties of ubiquitin-associated domains. Nat Struct Mol Biol 12, 708–714. https://doi.org/10.1038/nsmb962

Reynolds, A.D., Glanzer, J.G., Kadiu, I., Ricardo-Dukelow, M., Chaudhuri, A., Ciborowski, P., Cerny, R., Gelman, B., Thomas, M.P., Mosley, R.L., Gendelman, H.E., 2008a. Nitrated alpha-synuclein-activated microglial profiling for Parkinson’s disease: Synuclein-induced microglia activation. Journal of Neurochemistry 104, 1504–1525. https://doi.org/10.1111/j.1471-4159.2007.05087.x

Reynolds, A.D., Kadiu, I., Garg, S.K., Glanzer, J.G., Nordgren, T., Ciborowski, P., Banerjee, R., Gendelman, H.E., 2008b. Nitrated Alpha-Synuclein and Microglial Neuroregulatory Activities. J Neuroimmune Pharmacol 3, 59–74. https://doi.org/10.1007/s11481-008-9100-z

Reynolds, A.D., Stone, D.K., Mosley, R.L., Gendelman, H.E., 2009. Nitrated α-Synuclein-Induced Alterations in Microglial Immunity Are Regulated by CD4+ T Cell Subsets. The Journal of Immunology 182, 4137–4149. https://doi.org/10.4049/jimmunol.0803982

Rostami, J., Holmqvist, S., Lindström, V., Sigvardson, J., Westermark, G.T., Ingelsson, M., Bergström, J., Roybon, L., Erlandsson, A., 2017. Human Astrocytes Transfer Aggregated Alpha-Synuclein via Tunneling Nanotubes. J. Neurosci. 37, 11835–11853. https://doi.org/10.1523/JNEUROSCI.0983-17.2017

Shoji, M., Harigaya, Yasuo, Sasaki, A., Ueda, K., Ishiguro, K., Matsubara, E., Watanabe, M., Ikeda, M., Kanai, M., Tomidokoro, Y., Shizuka, M., Amari, M., Kosaka, K., Nakazato, Y., Okamoto, K., Hirai, S., 2000. Accumulation of NACP/alpha -synuclein in Lewy body disease and multiple system atrophy. Journal of Neurology, Neurosurgery & Psychiatry 68, 605–608. https://doi.org/10.1136/jnnp.68.5.605

Sofroniew, M.V., Vinters, H.V., 2010. Astrocytes: biology and pathology. Acta Neuropathologica 29.

Song, Y.J.C., Halliday, G.M., Holton, J.L., Lashley, T., O’Sullivan, S.S., McCann, H., Lees, A.J., Ozawa, T., Williams, D.R., Lockhart, P.J., Revesz, T.R., 2009. Degeneration in Different Parkinsonian Syndromes Relates to Astrocyte Type and Astrocyte Protein Expression. J Neuropathol Exp Neurol 68, 1073–1083. https://doi.org/10.1097/NEN.0b013e3181b66f1b

Sorrentino, Z.A., Goodwin, M.S., Riffe, C.J., Dhillon, J.-K.S., Xia, Y., Gorion, K.-M., Vijayaraghavan, N., McFarland, K.N., Golbe, L.I., Yachnis, A.T., Giasson, B.I., 2019. Unique α-synuclein pathology within the amygdala in Lewy body dementia: implications for disease initiation and progression. acta neuropathol commun 7, 142. https://doi.org/10.1186/s40478-019-0787-2

Spillantini, M.G., Crowther, R.A., Jakes, R., Hasegawa, M., Goedert, M., 1998. Alpha-synuclein in filamentous inclusions of Lewy bodies from Parkinson’s disease and dementia with Lewy bodies. Proceedings of the National Academy of Sciences 95, 6469–6473. https://doi.org/10.1073/pnas.95.11.6469

Takeda, A., Hashimoto, M., Mallory, M., Sundsumo, M., Hansen, L., Masliah, E., 2000. C-terminal α-synuclein immunoreactivity in structures other than Lewy bodies in neurodegenerative disorders. Acta Neuropathol 99, 296–304. https://doi.org/10.1007/PL00007441

Tanji, K., Mori, F., Mimura, J., Itoh, K., Kakita, A., Takahashi, H., Wakabayashi, K., 2010. Proteinase K-resistant α-synuclein is deposited in presynapses in human Lewy body disease and A53T α-synuclein transgenic mice. Acta Neuropathol 120, 145–154. https://doi.org/10.1007/s00401-010-0676-z

Terada, S., Ishizu, H., Haraguchi, T., Takehisa, Y., Tanabe, Y., Kawai, K., Kuroda, S., 2000. Tau-negative astrocytic star-like inclusions and coiled bodies in dementia with Lewy bodies. Acta Neuropathologica 100, 464–468. https://doi.org/10.1007/s004010000213

Terada, S., Ishizu, H., Yokota, O., Tsuchiya, K., Nakashima, H., Ishihara, T., Fujita, D., Uéda, K., Ikeda, K., Kuroda, S., 2003. Glial involvement in diffuse Lewy body disease. Acta Neuropathol 105, 163–169. https://doi.org/10.1007/s00401-002-0622-9

Thomas, M.P., Chartrand, K., Reynolds, A., Vitvitsky, V., Banerjee, R., Gendelman, H.E., 2007. Ion channel blockade attenuates aggregated alpha synuclein induction of microglial reactive oxygen species: relevance for the pathogenesis of Parkinson’s disease. J Neurochem 100, 503–519. https://doi.org/10.1111/j.1471-4159.2006.04315.x

Vaikath, N.N., Majbour, N.K., Paleologou, K.E., Ardah, M.T., van Dam, E., van de Berg, W.D.J., Forrest, S.L., Parkkinen, L., Gai, W.-P., Hattori, N., Takanashi, M., Lee, S.-J., Mann, D.M.A., Imai, Y., Halliday, G.M., Li, J.-Yi., El-Agnaf, O.M.A., 2015. Generation and characterization of novel conformation-specific monoclonal antibodies for α-synuclein pathology. Neurobiology of Disease 79, 81–99. https://doi.org/10.1016/j.nbd.2015.04.009

Volpicelli-Daley, L.A., Luk, K.C., Patel, T.P., Tanik, S.A., Riddle, D.M., Stieber, A., Meaney, D.F., Trojanowski, J.Q., Lee, V.M.-Y., 2011. Exogenous α-Synuclein Fibrils Induce Lewy Body Pathology Leading to Synaptic Dysfunction and Neuron Death. Neuron 72, 57–71. https://doi.org/10.1016/j.neuron.2011.08.033

Wakabayashi, K., Hayashi, S., Yoshimoto, M., Kudo, H., Takahashi, H., 2000. NACP/α-synuclein-positive filamentous inclusions in astrocytes and oligodendrocytes of Parkinson’s disease brains. Acta Neuropathol 99, 14–20. https://doi.org/10.1007/PL00007400

Wilhelmsson, U., Bushong, E.A., Price, D.L., Smarr, B.L., Phung, V., Terada, M., Ellisman, M.H., Pekny, M., 2006. Redefining the concept of reactive astrocytes as cells that remain within their unique domains upon reaction to injury. Proceedings of the National Academy of Sciences 103, 17513–17518. https://doi.org/10.1073/pnas.0602841103

Yamanaka, K., Chun, S.J., Boillee, S., Fujimori-Tonou, N., Yamashita, H., Gutmann, D.H., Takahashi, R., Misawa, H., Cleveland, D.W., 2008. Astrocytes as determinants of disease progression in inherited amyotrophic lateral sclerosis. Nat Neurosci 11, 251–253. https://doi.org/10.1038/nn2047

Zhang, Wei, Wang, T., Pei, Z., Miller, D.S., Wu, X., Block, M.L., Wilson, B., Zhang, Wanqin, Zhou, Y., Hong, J.-S., Zhang, J., 2005. Aggregated α-synuclein activates microglia: a process leading to disease progression in Parkinson’s disease. The FASEB Journal 19, 533–542. https://doi.org/10.1096/fj.04-2751com

