## Supplementary Figures and Tables for "Prominent astrocytic alpha-synuclein pathology with unique post-translational modification signatures unveiled across Lewy body disorders"

### SUPPLEMENTARY FIGURE 1

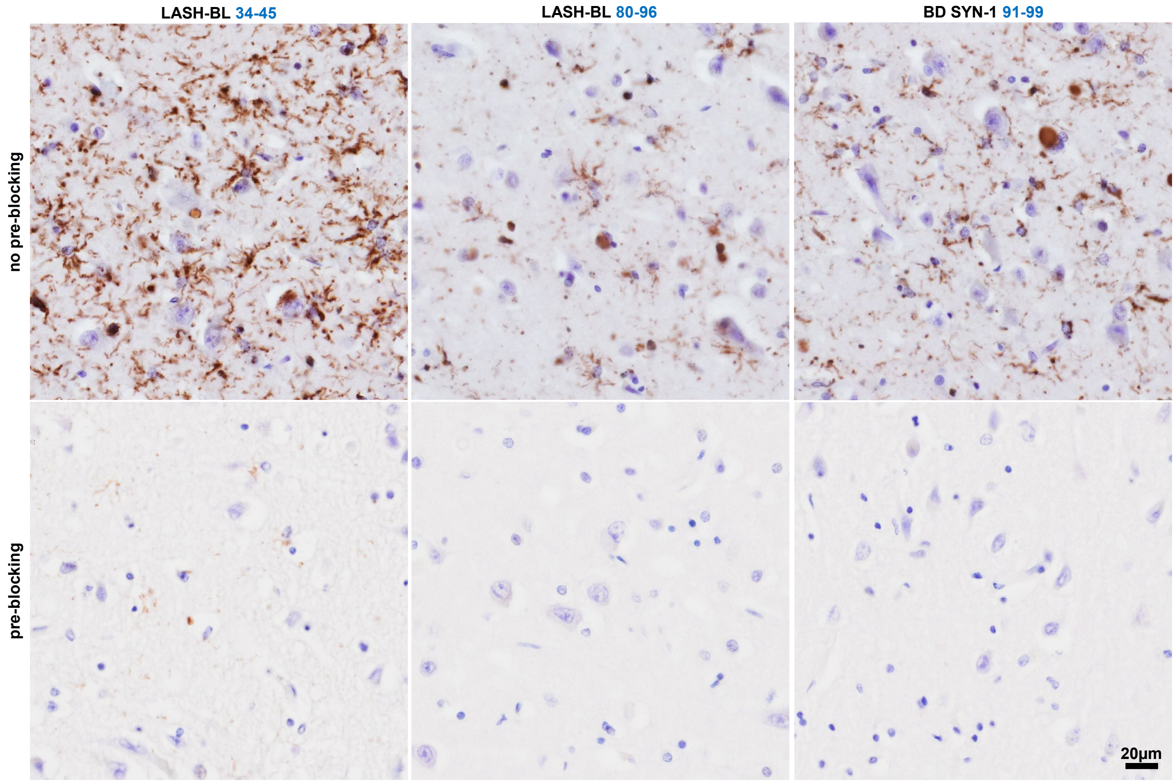

#### SUPPLEMENTARY FIGURE 2

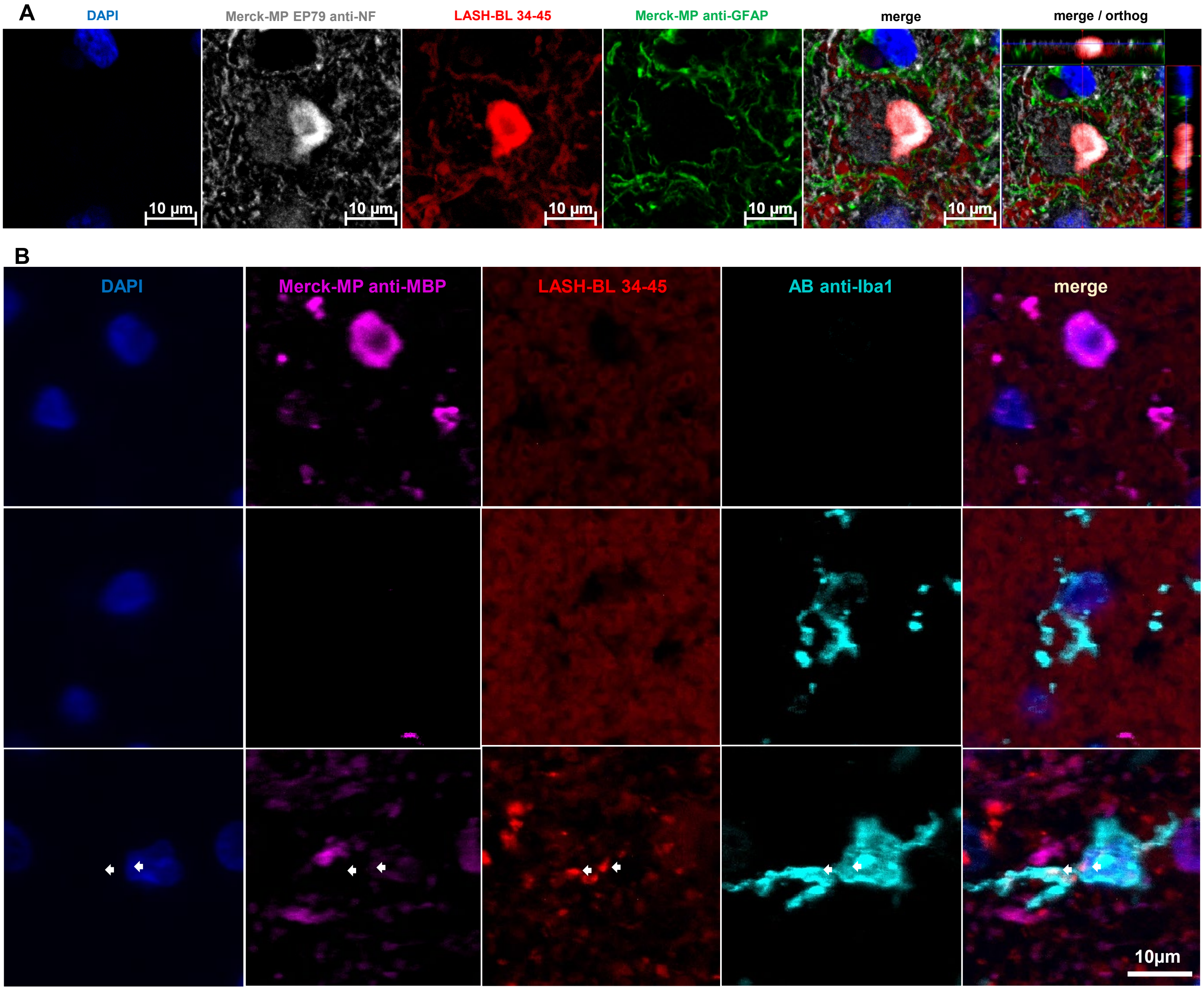

### SUPPLEMENTARY FIGURE 3

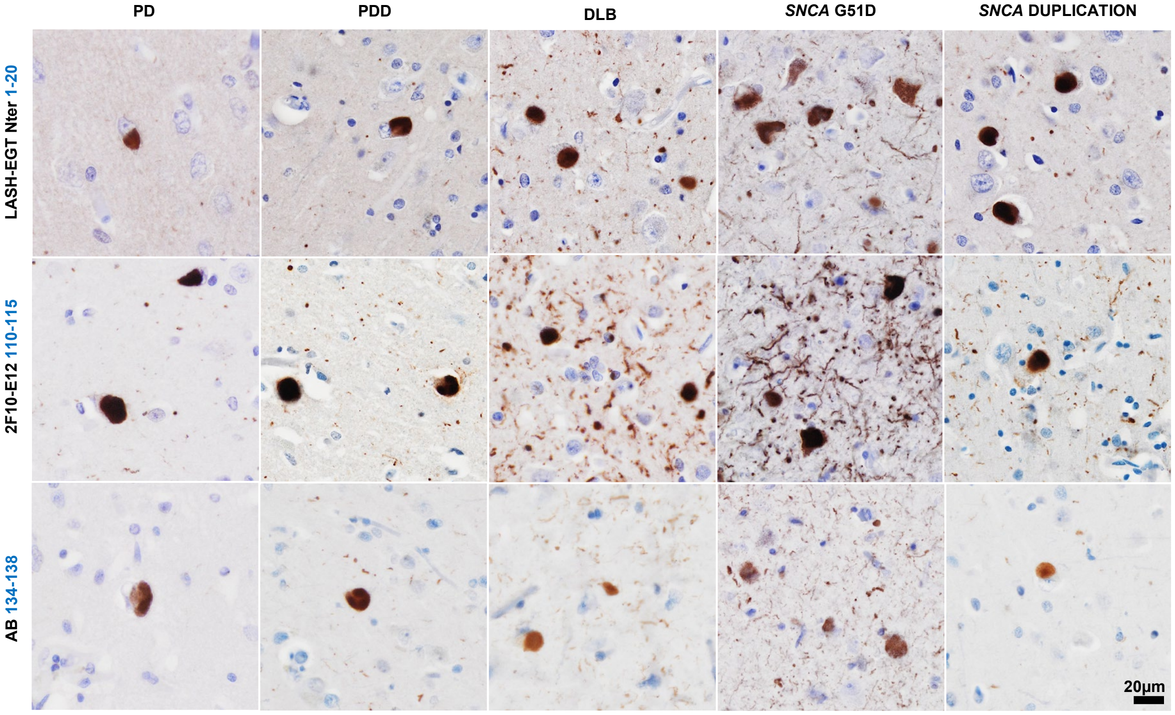

#### SUPPLEMENTARY FIGURE 4

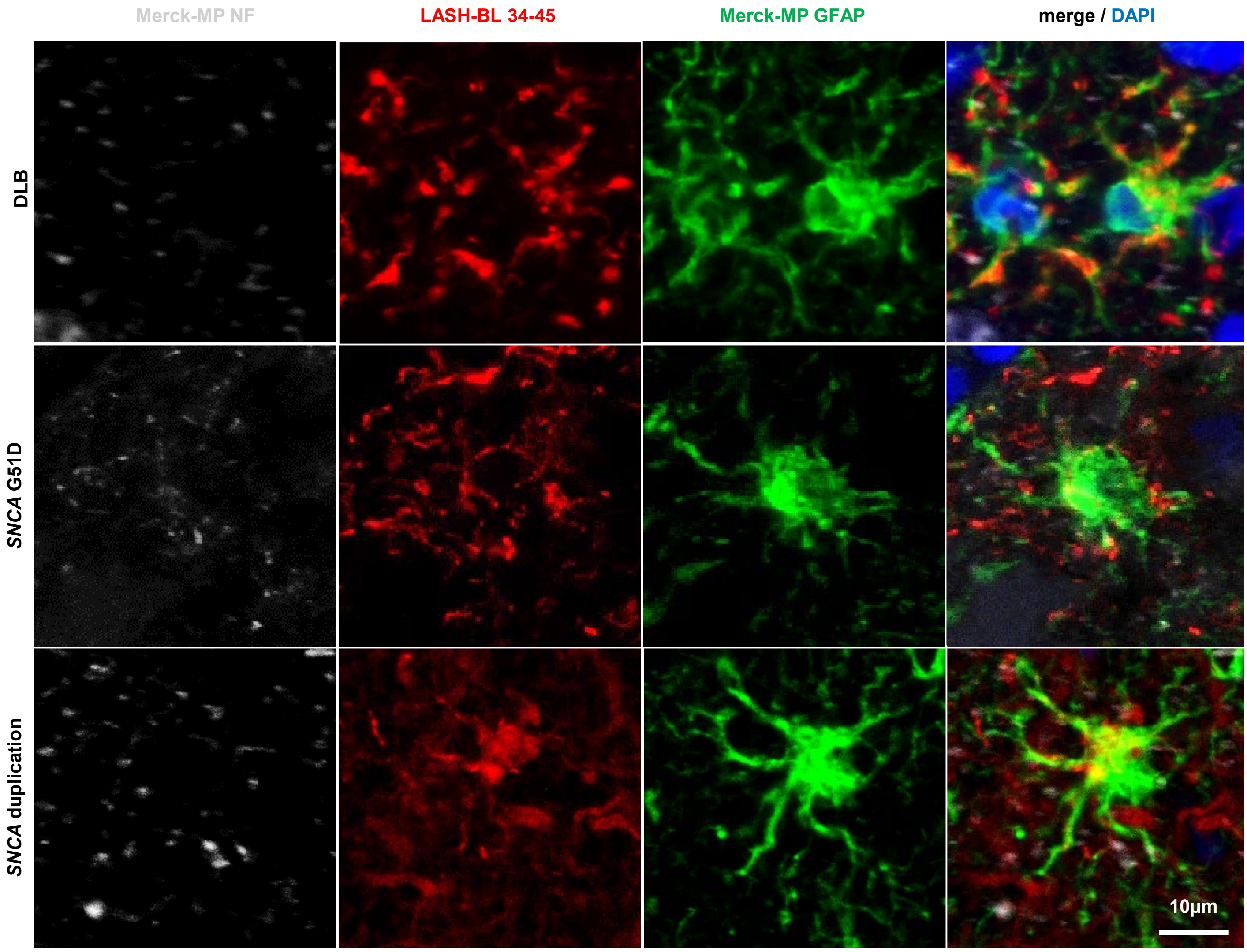

#### SUPPLEMENTARY FIGURE 5

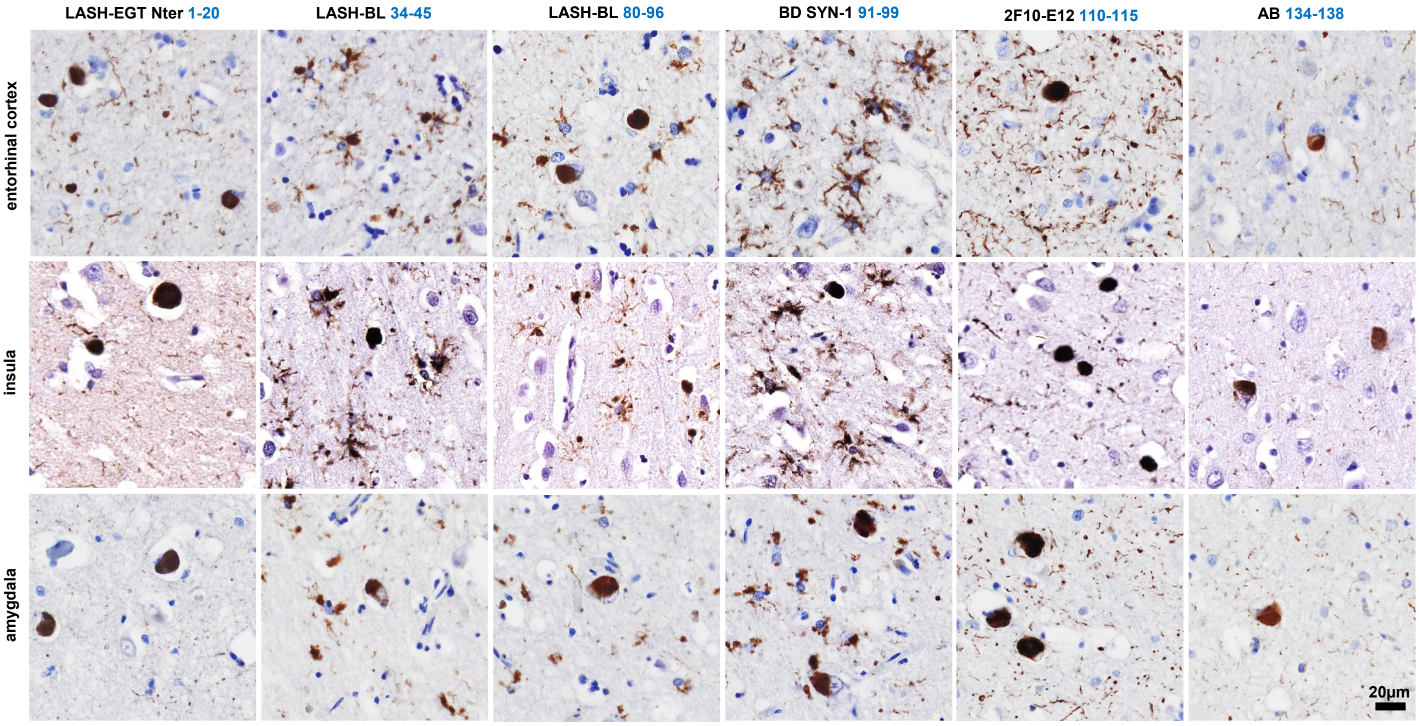

**Supplementary Table 1:** The primary and secondary antibodies included in this study.

| primary antibodies | | | | | | | |
|---|---|---|---|---|---|---|---|
| **antibody** | **epitope** | **species/ clonality** | **company / catalogue #** | **IHC: antigen retrieval** | **IHC: dilution** | **IF: dilution*** | **SB: dilution** |
| LASH-EGTNter | aSyn 1-20 | rab pc | na | AC+FA | 1:15,000 | na | 1:1,000 |
| LASH-BL 34-45 | aSyn 34-45 | mus mc | Biolegend #849101 | FA | 1:30,000 | 1:10,000 | na |
| LASH-BL 80-96 | aSyn 80-96 | mus mc | Biolegend #848302 | FA | 1:20,000 | na | na |
| BD SYN-1 | aSyn 91-99 | mus mc | BD #BD610787 | FA | 1:5,000 | na | 1:2,000 |
| BL 4B12 | aSyn 103-108 | mus mc | Biolegend #807801 | AC+FA | 1:100,000 | na | na |
| 2F10-E12 | aSyn 110-115 | mus mc | na | AC+FA | 1:10,000 | na | na |
| AB 134-138 | aSyn 134-138 | rab pc | Abcam #ab131508 | AC+FA | 1:25,000 | na | na |
| LASH-EGT pY39 | aSyn pY39 | rab pc | na | AC+FA | 1:500 | na | na |
| LASH-BL pY39 | aSyn pY39 | mus mc | Biolegend #849201 | AC+FA | 1:2,000 | 1:500 | 1:1,000 |
| LASH pS87 | aSyn pS87 | rab pc | na | FA | 1:600 | na | na |
| AB pY125 | aSyn pY125 | rab pc | Abcam #ab10789 | FA | 1:500 | na | na |
| AB EP1536Y | aSyn pS129 | rab mc | Abcam #ab51253 | AC+FA | 1:60,000 | 1:10,000 | na |
| AB pY133 | aSyn pY133 | rab pc | Abcam #ab194910 | AC+FA | 1:400 | na | na |
| AB pY136 | aSyn pY136 | rab pc | Abcam #ab131491 | FA | 1:100 | na | na |
| LASH-EGT nY39 | aSyn nY39 | rab pc | na | FA | 1:1,000 | 1:500 | 1:200 |
| 6A3-E9 | aSyn-120 | mus mc | na | AC+FA | 1:2,500 | na | na |
| 5G4 | 44-57 // agg-aSyn | mus mc | Merck-Millipore #MABN389 | AC+FA | 1:5,000 | na | na |
| SYNO4 | agg-aSyn | mus mc | na | FA | 1:5,000 | na | na |
| Merck-MP anti-GFAP | GFAP | ck pc | Merck-Millipore #AB5541 | na | na | 1:500 | na |
| Merck-MP EP79 anti-NF | NF | rab mc | Merck-Millipore #302R-1 | na | na | 1:500 | na |
| AB anti-Iba1 | Iba1 | goat pc | Abcam #ab5076 | na | na | 1:80 | na |
| Merck-MP anti-MBP | MBP | rab pc | Merck-Millipore #AB5864 | na | na | 1:2,000 | na |
| AB EPR8830 recomb anti-Ub | ubiquitin | rab mc | Abcam #ab134953 | na | na | 1:800 | na |
| Proteintech anti-p62 | p62 | rab pc | Proteintech #18420-1-AP | na | na | 1:200 | na |
| *AC+FA pre-treatment was applied for IF studies for all antibodies. | | | | | | | |

| secondary antibodies | | | | |
|---|---|---|---|---|
| **antibody** | **dilution** | **company** | **catalogue #** | **application** |
| donkey anti-chicken 488 | 1:400 | Jackson ImmunoResearch | 703-545-155 | IF |
| goat anti-chicken 568 | 1:400 | ThermoFisher | A-11041 | IF |
| donkey anti-goat 488 | 1:1,000 | ThermoFisher | A-11055 | IF |
| donkey anti-rabbit 568 | 1:1,000 | ThermoFisher | A-10042 | IF |
| donkey anti-mouse 568 | 1:1,000 | ThermoFisher | A-10037 | IF |
| donkey anti-mouse 647 | 1:1,000 | ThermoFisher | A-31571 | IF |
| IRDye goat anti-mouse 680 | 1:20,000 | Li-Cor | 926-68070 | SB |
| IRDye goat anti-rabbit 800 | 1:20,000 | Li-Cor | 926-32211 | SB |

AC = autoclave; agg-aSyn = aggregated alpha-synucleina; aSyn = alpha-synuclein; FA = formic acid; GFAP = glial fibrillary acidic protein; Iba1 = ionised calcium binding adaptor protein 1; IF = immunofluorescence; IHC = immunohistochemistry; MBP = myelin basic protein; mc = monoclonal; mus = mouse; na = not applicable; NF = neurofilament; pc = polyclonal; rab = rabbit; SB = slot blot
